# Nucleo-cytoplasmic shuttling of splicing factor SRSF1 is required for development and cilia function

**DOI:** 10.1101/2020.09.04.263251

**Authors:** Fiona Haward, Magdalena M. Maslon, Patricia L. Yeyati, Nicolas Bellora, Jan N. Hansen, Stuart Aitken, Jennifer Lawson, Alex von Kriegsheim, Dagmar Wachten, Pleasantine Mill, Ian R. Adams, Javier F. Cáceres

## Abstract

Shuttling RNA-binding proteins coordinate nuclear and cytoplasmic steps of gene expression. The SR family proteins regulate RNA splicing in the nucleus and a subset of them, including SRSF1, shuttles between the nucleus and cytoplasm affecting post-splicing processes. However, the physiological significance of this remains unclear. Here, we used genome editing to knock-in a nuclear retention signal (NRS) in *Srsf1* to create a mouse model harboring an SRSF1 protein that is retained exclusively in the nucleus. *Srsf1^NRS/NRS^* mutants displayed small body size, hydrocephalus and immotile sperm, all traits associated with ciliary defects. We observed reduced translation of a subset of mRNAs and decreased abundance of proteins involved in multiciliogenesis, with disruption of ciliary ultrastructure and motility in cells derived from this mouse model. These results demonstrate that SRSF1 shuttling is used to reprogram gene expression networks in the context of high cellular demands, as observed here, during motile ciliogenesis.

## Introduction

Alternative splicing (AS) is an essential step in the gene expression cascade that generates a vast number of mRNA isoforms to shape the proteomes of multicellular eukaryotes (Baralle and Giudice, 2017; Nilsen and Graveley, 2010; Ule and Blencowe, 2019). It is largely controlled by the binding of RNA-binding proteins (RBPs) in a manner dependent on the cellular context (Fu and Ares, 2014). Additional layers of regulation include alterations in chromatin state and the co-transcriptional nature of the splicing process, including the rate of RNA Polymerase II (RNAPII) elongation (Maslon et al., 2019; Naftelberg et al., 2015; Saldi et al., 2016). Splicing alterations are found in human disease, and are particularly common in cancer due to mutations in cis-acting splicing-regulatory elements or in components of the splicing machinery (Anczukow et al., 2016; Bonnal et al., 2020; Zhang and Manley, 2013).

The serine/arginine-rich (SR) family proteins are among the most extensively characterized regulators of pre-mRNA splicing (Wegener and Müller-McNicoll, 2019; Zhou and Fu, 2013). Their modular domain structure comprises one or two RNA recognition motifs (RRMs) at their N-termini that contribute to sequence-specific RNA-binding and a C-terminal RS domain (arginine and serine repeats) that promotes protein-protein interactions (Howard and Sanford, 2015). The RS domain also acts as a nuclear localization signal by promoting the interaction with the SR protein nuclear import receptor, transportin-SR (Cáceres et al., 1997; Kataoka et al., 1999; Lai et al., 2000). The role of SR proteins in alternative splicing regulation can be antagonized by heterogeneous nuclear ribonucleoproteins (hnRNPs). Thus, the relative ratio of these two protein families can regulate alternative splicing patterns that generate specific cell lineages and tissues during development (Busch and Hertel, 2012; Hanamura et al., 1998; Zhu et al., 2001).

Besides the clearly defined nuclear roles of SR family proteins, a subset of these, of which SRSF1 is the prototype, shuttle continuously from the nucleus to the cytoplasm and are involved in post-splicing activities (Cáceres et al., 1998; Cowper et al., 2001; Sapra et al., 2009). These include mRNA export as well as cytoplasmic roles such as mRNA translation, nonsense-mediated decay (NMD) and regulation of RNA stability (reviewed by (Long and Caceres, 2009; Twyffels et al., 2011)). Several protein kinases phosphorylate the RS domain of SR proteins, including the SRPK family (Gui et al., 1994) and the Clk/Sty family dual-specificity kinases (Prasad et al., 1999). Whereas phosphorylated SR proteins are required for spliceosome assembly, subsequent dephosphorylation is required for splicing catalysis and for sorting shuttling and non-shuttling SR proteins in the nucleus (Lin et al., 2005). Our previous work demonstrated that hypophosphorylated SRSF1 is associated with polyribosomes and promotes translation of mRNAs in an mTOR-dependent manner (Michlewski et al., 2008; Sanford et al., 2005, 2004). We subsequently showed that translational targets of SRSF1 predominantly encode proteins that localize to centrosomes and are required for cell cycle regulation and chromosome segregation (Maslon et al., 2014). The shuttling ability of individual SR proteins has also been usurped by viruses, as seen with the role of individual SR proteins in promoting translation of viral transcripts, including poliovirus, MMPV and HIV (Bedard et al., 2007; Swartz et al., 2007).

SRSF1 is essential for cellular viability (Lin et al., 2005; Wang et al., 1996) and has been shown to act as an oncogene and to promote mammary epithelial cell transformation (Anczuków et al., 2012; Karni et al., 2007). In living animals, its targeted disruption results in early embryonic lethality that cannot be rescued by other SR proteins (Möröy and Heyd, 2007; Xu et al., 2005). Furthermore, a T cell-restricted *Srsf1*-deficient mice develops systemic autoimmunity and lupus-nephritis (Katsuyama et al., 2019; Paz et al., 2020), whereas its deletion in myogenic progenitors leads to defects in neuromuscular junctions (Liu et al., 2020). This evidence highlights the splicing roles of SRSF1 in cellular transformation or in the development of tissues. By contrast, the physiological relevance of its post-splicing activities has remained fully enigmatic. In particular, it is currently not clear to what extent SRSF1 might regulate mRNA translation in more physiological contexts, or what the physiological relevance of this regulation might be.

Here, we have engineered a mouse model of a non-shuttling endogenous SRSF1 protein that is exclusively retained in the nucleus. We show that post-splicing activities of SRSF1 in the cytoplasm are dispensable for embryonic development; however, mutant animals display characteristic motile cilia phenotypes. We also observed that the lack of cytoplasmic SRSF1 leads to reduced translation of a subset of mRNAs in mouse neural stem cells (NSCs) and a decrease in the abundance of proteins involved in multiciliogenesis in tracheal cultures. We conclude that nucleo-cytoplasmic shuttling of SRSF1 is indeed required for proper development and primarily affects mRNA translation, contributing to the biogenesis and function of motile cilia.

## Results

### Generation of a non-shuttling SRSF1 protein mouse model

To create an *in vivo* mouse model expressing only a nuclear-retained SRSF1 protein, we inserted a potent nuclear retention signal (NRS) at the C-terminus of the *Srsf1* genomic locus (***Figure 1A***). This sequence is naturally present in the non-shuttling SR protein SRSF2 and when fused to SRSF1 prevents its shuttling when overexpressed (Cazalla et al., 2002; Maslon et al., 2014; Sanford et al., 2004). We designed a CRISPR/Cas9-assisted strategy in which an NRS sequence, a small linker and a T7 tag were introduced at the C-terminus of the canonical SRSF1 isoform (Sun et al., 2010) (***Figure 1B***, ***Figure 1-figure supplement 1***). This approach minimizes perturbation of the N-terminal RRM domains of SRSF1 that are crucial for RNA recognition and its function in splicing (Cléry et al., 2013) The presence of the T7 tag facilitates both visualization of the tagged protein (referred herein as SRSF1-NRS), as well as biochemical experiments. In contrast to the completely penetrant early embryonic lethality of *Srsf1^−/−^* knockout mice (Xu et al., 2005), *Srsf1^NRS/NRS^* mice were born from *Srsf1^+/NRS^* intercrosses and survived postnatally to the point of genotyping (day 14) (***Figure 1C***). However, postnatal *Srsf1^NRS/NRS^* mice displayed overt phenotypic abnormalities from this stage onwards (see later). These findings indicate that the cytoplasmic functions for SRSF1 allele are mostly dispensable for embryonic development. We confirmed that the SRSF1-NRS fusion protein is expressed in liver tissue and embryonic stem cells (ESCs) derived from homozygous *Srsf1^NRS/NRS^* mice (***Figure 1D***).

**Figure 1.**
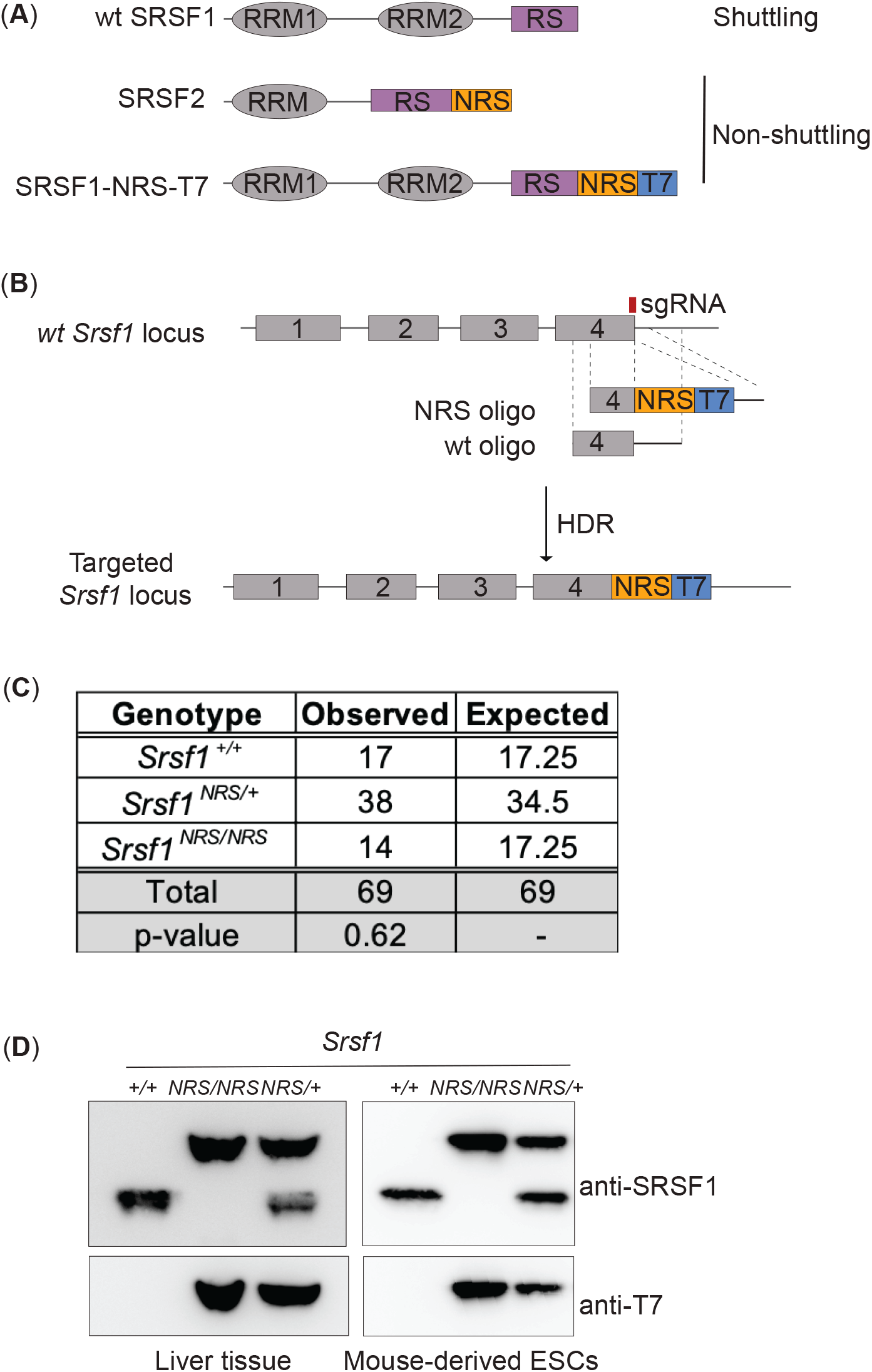
Generation of a nuclear restrained SRSF1 (*Srsf1^NRS/NRS^*) knock-in mouse model. (**A**) Domain structure and nucleo-cytoplasmic shuttling ability of wild-type (WT) SRSF1 and SRSF2 proteins, and the artificial fusion protein SRSF1-NRS, used in this study. (**B**) Schematic representation of the *Srsf1* locus and the CRISPR/Cas9 strategy used to introduce an NRS-T7 sequence at its C-terminus. The nucleotide sequence of the introduced NRS is identical to that present in the endogenous mouse *Srsf2* gene (Cazalla et al., 2002). Genotyping results are shown in ***Figure 1—figure supplement 1**.* (**C**) *Srsf1^NRS/NRS^* homozygous knock-in mice complete embryogenesis and are viable postnatally. The number of pups obtained from *Srsf1^+/NRS^* intercrosses with indicated genotypes at postnatal day 14 are indicated. The expected Mendelian numbers and χ^2^-test P value (P=0.62, n=69) are also shown. (**D**) Expression of the SRSF1-NRS protein from mouse liver tissue of three targeted mice (P14) (left panel) and in three ES cell lines derived from *Srsf1^NRS/NRS^* mice (right panel). In both panels, SRSF1 and T7 antibodies were used for Western blot analysis to detect the knock-in protein.

We and others have previously shown that the SRSF1-NRS chimeric protein is retained in the nucleus (Cazalla et al., 2002; Lin et al., 2005). Here, we analyzed the shuttling ability of endogenous SRSF1-NRS protein in neural stem cells (NSCs) differentiated *in vitro* from ESCs derived from *Srsf1^NRS/NRS^* animals, using an inter-species heterokaryon assay (Piñol-Roma and Dreyfuss, 1992). As expected, we observed that SRSF1-NRS protein was only detected in the mouse but not in human nuclei, confirming that it is indeed restricted to the mouse nucleus (***Figure 1-figure supplement 2***, lower panel). As a positive control, we transiently expressed a T7-tagged WT-SRSF1 in mouse NSCs and clearly observed its presence in the recipient HeLa nuclei, underlining the innate shuttling activity of WT SRSF1 (***Figure 1-figure supplement 2***, upper panel). This confirms that our targeting strategy was successful and the resulting SRSF1-NRS fusion protein expressed from the endogenous *Srsf1* locus is in fact restricted to the nucleus.

### Srsf1^NRS/NRS^ mice are growth restricted

The viability of *Srsf1^NRS/NRS^* embryos contrasts with the reported lethality of *Srsf1* null embryos by E7.5 (Xu et al., 2005). Strikingly, homozygous *Srsf1^NRS/NRS^* mice displayed numerous severe post-natal phenotypes. First, these knock-in mice were visibly smaller in size, including those that survived up to 7.5-months of age, being on average 30% lighter in weight than littermate controls (Student’s t-test, p<0.0001), suggestive of a growth restriction phenotype (***Figure 2A***). To investigate whether the growth restriction observed in *Srsf1^NRS/NRS^* mice arises during embryogenesis, E12.5 embryos and placentas were harvested from four *Srsf1^+/NRS^* intercross litters. *Srsf1^NRS/NRS^* embryos were grossly phenotypically normal at this stage of development (***Figure 2B***), with no difference in embryo or placenta weights compared to their *Srsf1^+/+^* (***Figure 2C***). These results indicate that the observed growth restriction does not arise from impaired embryogenesis or abnormal maternal nutrient transfer across the placenta, suggesting that postnatal growth is specifically affected in *Srsf1^NRS/NRS^* animals.

**Figure 2.**
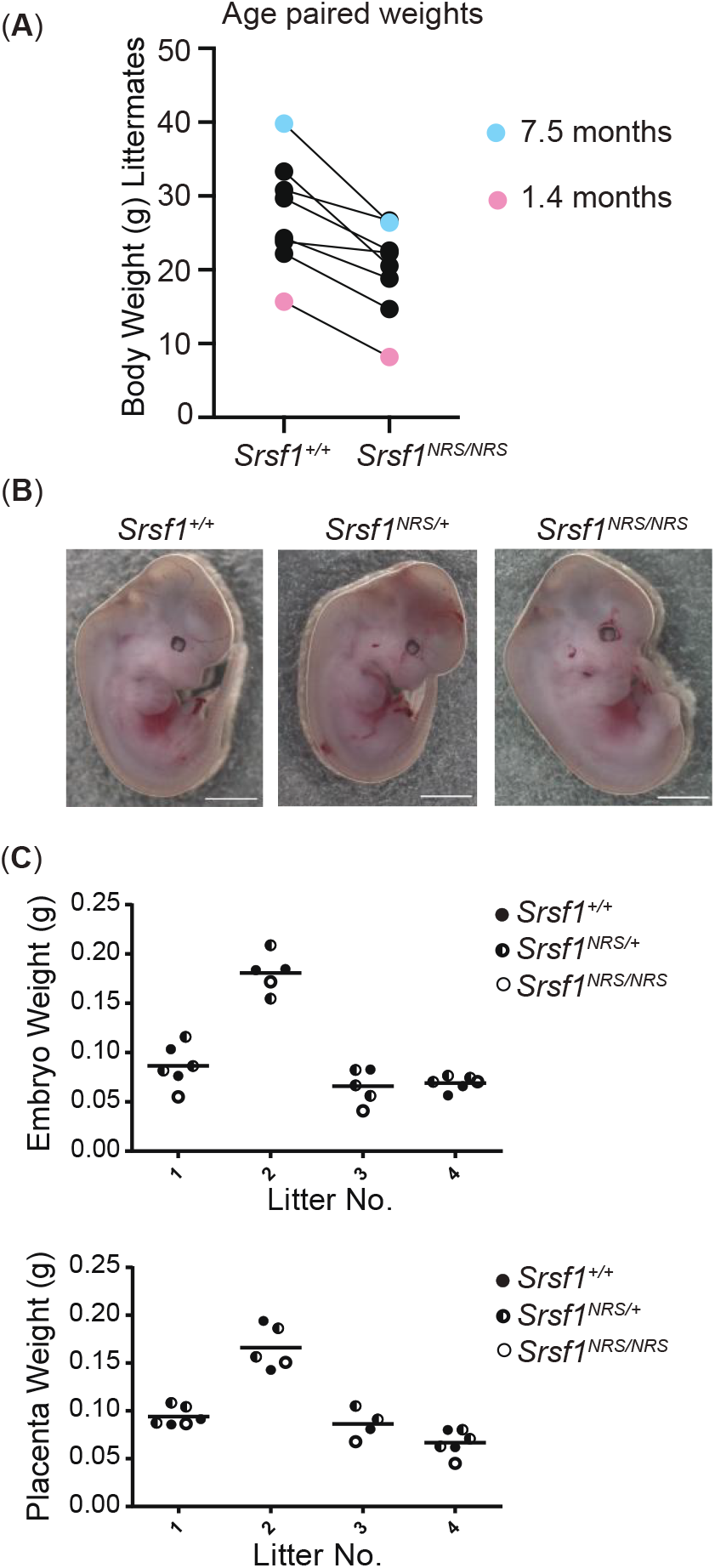
Homozygosity for the *Srsf1^NRS^* allele causes postnatal growth restriction. (**A**) Whole body weight of sex-matched littermate pairs indicated by connecting lines. Animals range from 1.4 (pink) to 7.5 months old (blue). (**B**) *Srsf1^NRS/NRS^* knock-in embryos at E13.5 are grossly phenotypically normal. *Srsf1^NRS/NRS^* embryos were represented in these litters at the expected Mendelian ratio (7 *Srsf1^+/+^*, 11 *Srsf1^+/NRS^*, 4 *Srsf1^NRS/NRS^*; χ^2^ P value=0.66). (**C**) Scatter plots showing the weight of whole embryos (top panel) and of placentas from four independent litters of E12.5 embryos (bottom panel).

### Srsf1^NRS/NRS^ mice perinatal phenotypes are indicative of defects in motile cilia

In addition to restricted growth, half of the *Srsf1^NRS/NRS^* animals developed hydrocephalus by P14. In contrast, no cases of hydrocephalus were observed in either the *Srsf1^+/+^* or *Srsf1^NRS/+^* littermates (***Figure 3A***). Hydrocephalus is caused by the accumulation of cerebrospinal fluid (CSF) resulting in dilatation of brain ventricles and increased pressure on the surrounding brain tissue, which can lead to cranial doming during the perinatal period (McAllister, 2012). As in humans, hydrocephalus can result in progressive brain enlargement, neurological dysfunction, seizures and death. Hydrocephalus can be caused by obstruction of the aqueducts or abnormal beating of cilia lining the brain ventricles thus preventing CSF flow (Ibañez-Tallon et al., 2004). However, we found no evidence of obstruction even in most severe cases of hydrocephalus (***Figure 3B***). *Srsf1^NRS/+^* animals were crossed onto a cilia cell cycle biosensor *ARL13B-Fucci2A* transgenic mice (Ford et al. 2018). Here, ubiquitous expression of the ciliary reporter ARL13B-Cerulean allows for live imaging of all ciliary types and revealed that ependymal cilia is present in *Srsf1^NRS/NRS^* animals (***Figure 3-figure supplement 1***).

**Figure 3.**
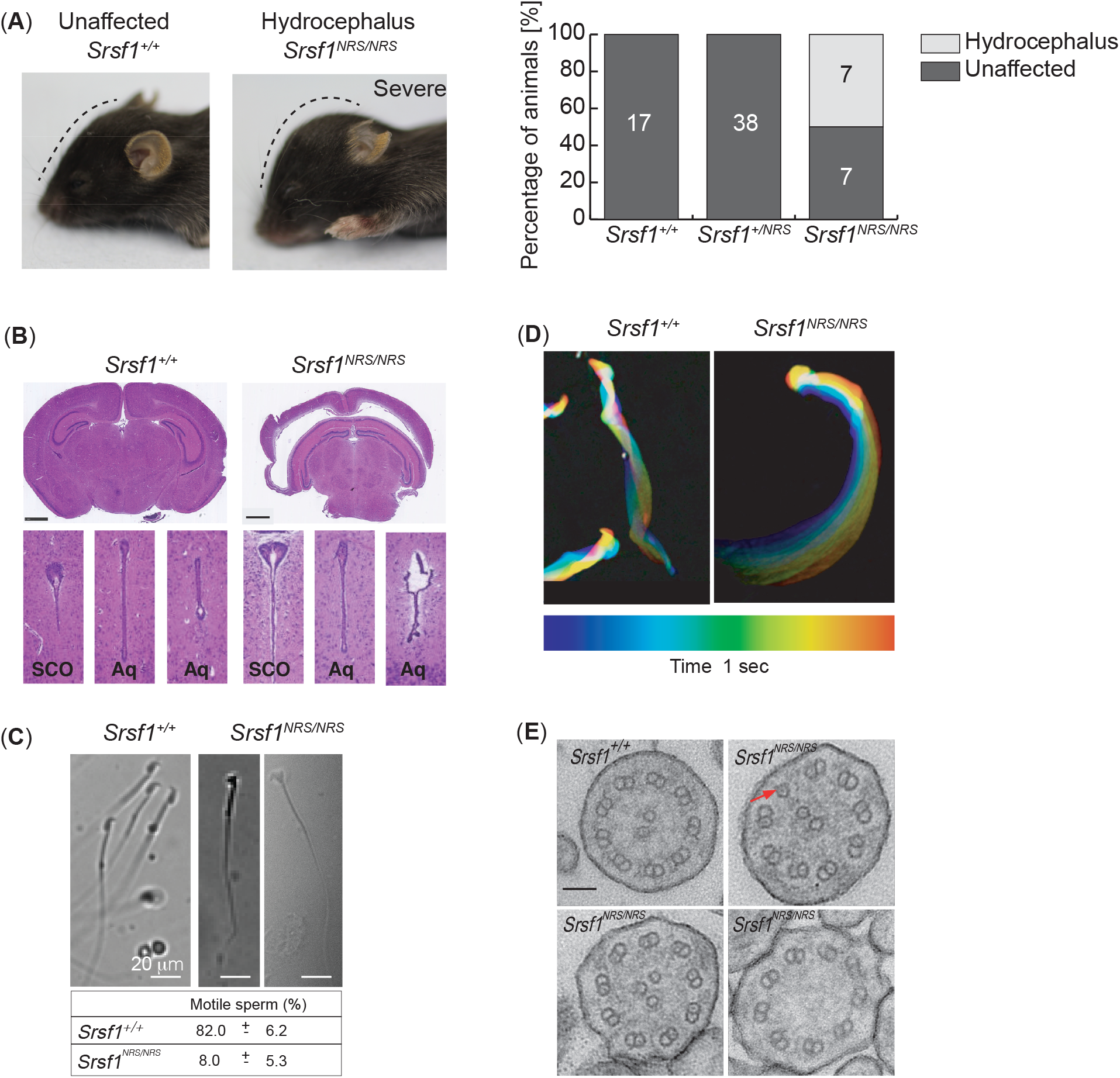
Homozygous *Srsf1^NRS/NRS^* mice develop hydrocephalus. (**A**) *Srsf1^NRS/NRS^* mice show signs of developing hydrocephalus (the curvature of the skull is depicted by a dashed line). Half of the mice culled from the first two cohorts developed externally visible hydrocephalus of a varying severity by P14, while hydrocephalus was not observed in *Srsf1^+/+^* or *Srsf1^NRS/+^* littermates. The barplot indicates the incidence of mice unaffected or with hydrocephalus (percentage and total number) in the 2 cohorts of mice used (p-value= 0.0013 and <0.0001, respectively; Fischer’s exact test). (**B**) H&E staining of coronal sections of brains at subcommissural organ (SCO) and aqueducts (Aq) of 3^rd^ and 4^th^ ventricles show no obvert stenosis. Scale bar: 1 mm. (**C**) Representative images illustrate abnormal head shape of some spermatozoa observed in *Srsf1^NRS/NRS^* littermates. Bottom panel shows percentage of motile sperm (N≥3 animals per genotype). (**D**) Color coding illustrates that the complex rotational pattern of *Srsf1^+/+^* spermatozoa, required to propel the sperm forward, is absent in the few motile *Srsf1^NRS/NRS^* spermatozoa. (**E**) Transmission electron microscopy of transverse sections of mTEC cilia showing (9+2) microtubules in *Srsf1^+/+^* or abnormal variations found in *Srsf1^NRS/NRS^*. Red arrow illustrates microtubule singlet. Scale bar: 100 nm.

In addition to hydrocephalus and growth restriction, *Srsf1^NRS/NRS^* males presented sperm with abnormal head morphology and a large proportion of immotile flagella (***Figure 3C)***, whereas those remaining motile exhibit abnormal waveforms (***Figure 3D***). Electron microscopy analysis also revealed ultrastructural defects in the motile ciliary axonemes of mouse tracheal epithelial cultures (mTECs) from *Srsf1^NRS/NRS^* mice; including lack of central pair (1/81), three central microtubules (10/81) or single outer microtubules (2/81) compared to control *Srsf1^+/+^* cultures (1 single outer microtubule out of n= 94) (Fig. 5G; *X^2^* of all microtubule defects < 0.001) ***(Figure 3E)***. These collective traits suggest that altered properties of motile cilia underlie the abnormal phenotypes of *Srsf1^NRS/NRS^* animals.

### The SRSF1-NRS protein does not affect mRNA splicing or global mRNA export

SRSF1 is essential for cellular and organism viability, which has been attributed to its splicing role in the nucleus (Lin et al., 2005; Xu et al., 2005). The viability of *Srsf1^NRS/NRS^* knock-in mice, strongly suggests that SRSF1-mediated splicing was not grossly affected in the homozygous animals. In addition, previous work using an exogenous SRSF1-NRS protein revealed that the nuclear splicing role of SRSF1 was essential and could be decoupled from its ability to shuttle (Lin et al 2005). We performed deep RNA-sequencing analysis on mouse embryonic fibroblasts (MEF) derived from *Srsf1^NRS/NRS^*, *Srsf1^NRS/+^* and *Srsf1^+/+^* littermates (***Figure 4-figure supplement 1A***, ***Figure 4-source data 1***). Splicing changes were analyzed using SUPPA2, a tool which displays differential splicing between conditions as changes in proportion spliced-in (ΔPSI) (Trincado et al., 2018). As such, SUPPA2 is advantageous as it permits analysis of multiple samples and accounts for biological variability between samples. This was important as our biological replicates were isolated from different embryos with the same genotype, that will have a certain degree of intrinsic heterogeneity. Pairwise comparisons of *Srsf1^+/+^*, *Srsf1^NRS/+^* and *Srsf1^NRS/NRS^* MEFs demonstrated that there were no gross splicing changes induced by the presence of the NRS insertion (***Figure 4-figure supplement 1A***, ***Figure 4-source data 1***). Out of 65318 transcripts analyzed, we observed only 26 changes in splicing considering ΔPSI =>0.2 (p<=0.01), which is comparable to the number of changes between *Srsf1^NRS/+^* and *Srsf1^+/+^* MEFs. Considering that *Srsf1^NRS/+^* mice do not have a quantifiable phenotype, the changes we observe likely represent stochastic changes in alternative splicing that could be explained by environmental or sampling factors. This confirms that precluding the shuttling ability of SRSF1 preserves its splicing function.

Shuttling SR proteins, including SRSF1, have been also proposed to function in nuclear mRNA export, through interactions with the export factor TAP/NXF1 (Hargous et al., 2006; Huang et al., 2003; Huang and Steitz, 2001). We performed high-throughput RNA sequencing analysis on cytoplasmic fractions from MEFs derived from *Srsf1^+/+^* or *Srsf1^NRS/NRS^* animals or NSCs differentiated *in vitro* from ESCs derived from *Srsf1^+/+^* or *Srsf1^NRS/NRS^* animals (***Figure 4-figure supplement 1B***, ***Figure 4-source data 2***). We found that the abundance of 121 and 57 mRNAs was decreased in the cytoplasm of *Srsf1^NRS/NRS^* MEFs and NSCs, respectively. In previous work, 225 transcripts were identified as the direct export targets of SRSF1 in P19 cells (Müller-McNicoll et al., 2016), however we found no overlap between those targets and mRNAs that change in this study. Similarly, cell fractionation followed by RT-qPCR of selected mRNAs in *Srsf1^+/+^* or *Srsf1^NRS/NRS^* cultures, revealed that the NRS insertion affected cytoplasmic levels of very few export candidates tested (***Figure 4-figure supplement 1C***). Taken together, our data shows that a nuclear restricted SRSF1 does not compromise pre-mRNA splicing and is associated with relatively few changes in mRNA export.

### Reduced translation of a subset of mRNAs in Srsf1^NRS/NRS^ NSCs

To determine whether the presence of a nuclear-retained SRSF1 protein affects mRNA translation *in vivo*, we performed polysomal shift analyses in different cell lines derived from *Srsf1^+/+^* and *Srsf1^NRS/NRS^* animals. We selected ESCs as a pluripotent cell type, NSCs to explore changes upon differentiation into cellular types relevant to the observed phenotypes of *Srsf1^NRS/NRS^* animals, and MEFs as a primary cell type. Cytoplasmic RNA was harvested from three biological replicates and RNA-seq and polysomal profiling was carried out to measure the transcriptome and translatome, respectively (***Figure 4A***, ***Figure 4-source data 3***). We confirmed the expression of the pluripotency gene *Oct4* (*Pou5f1*) in ESCs, whereas the co-expression of the neural stem cell markers *Nestin* (*Nes*) and *Sox2* in NSCs, indicated that the neural lineage has been successfully induced (***Figure 4-source data 2***). Finally, *Thy1* was chosen as a fibroblast marker for MEF cultures. Cytoplasmic extracts were fractionated across 10-45% sucrose gradients and RNA isolated from subpolysomal and heavy polysomal fractions, followed by high throughput sequencing, as previously described (Maslon et al., 2014). To accurately identify those mRNAs whose translation is responsive to the presence of cytoplasmic SRSF1 and remain associated with ribosomes during the fractionation procedure, we calculated polysome indices (PI) for each expressed gene, normalized to transcripts per million (TPM) and the polysome shift ratio (PSR) (PSR = log2(PI_SRSF1-NRS/PI_SRSF1)) between *Srsf1^+/+^* and *Srsf1^NRS/NRS^* cultures (see Methods). We used these values to determine transcripts that were significantly depleted (p<0.05) from polysomes in *Srsf1^NRS/NRS^* cells, as translation of such transcripts is likely to be directly compromised by the lack of shuttling SRSF1. This analysis revealed substantial changes in the association of mRNAs with polysomes in *Srsf1^NRS/NRS^* NSCs and MEFs, with a much lower proportion of changes in ESCs (***Figure 4B***). We found that 13 genes (total of 88) in ESCs, 1077 genes (total of 1258) in NSCs and 464 genes (total of 733) in MEFs were under-represented in the polysomal fractions of *Srsf1^NRS/NRS^* samples, strongly suggesting that SRSF1 is directly involved in their translation. When we compared translational changes considering only genes commonly expressed in all three cell types (9073 in total), we observed that this had marginally affected the number of genes being affected in all three cell types (***Figure 4-figure supplement 2***). Therefore, the observed translational changes are not restricted to lineage specific transcripts but instead seem to correlate with ongoing differentiation programs or metabolic demands of each cell type. These results suggest that differentiating cells like NSCs and to a lesser extent MEFs, require SRSF1 for the translation of a subset of mRNAs. Interestingly, these SRSF1-dependent translational functions are dispensable in pluripotent cells like ESCs.

**Figure 4.**
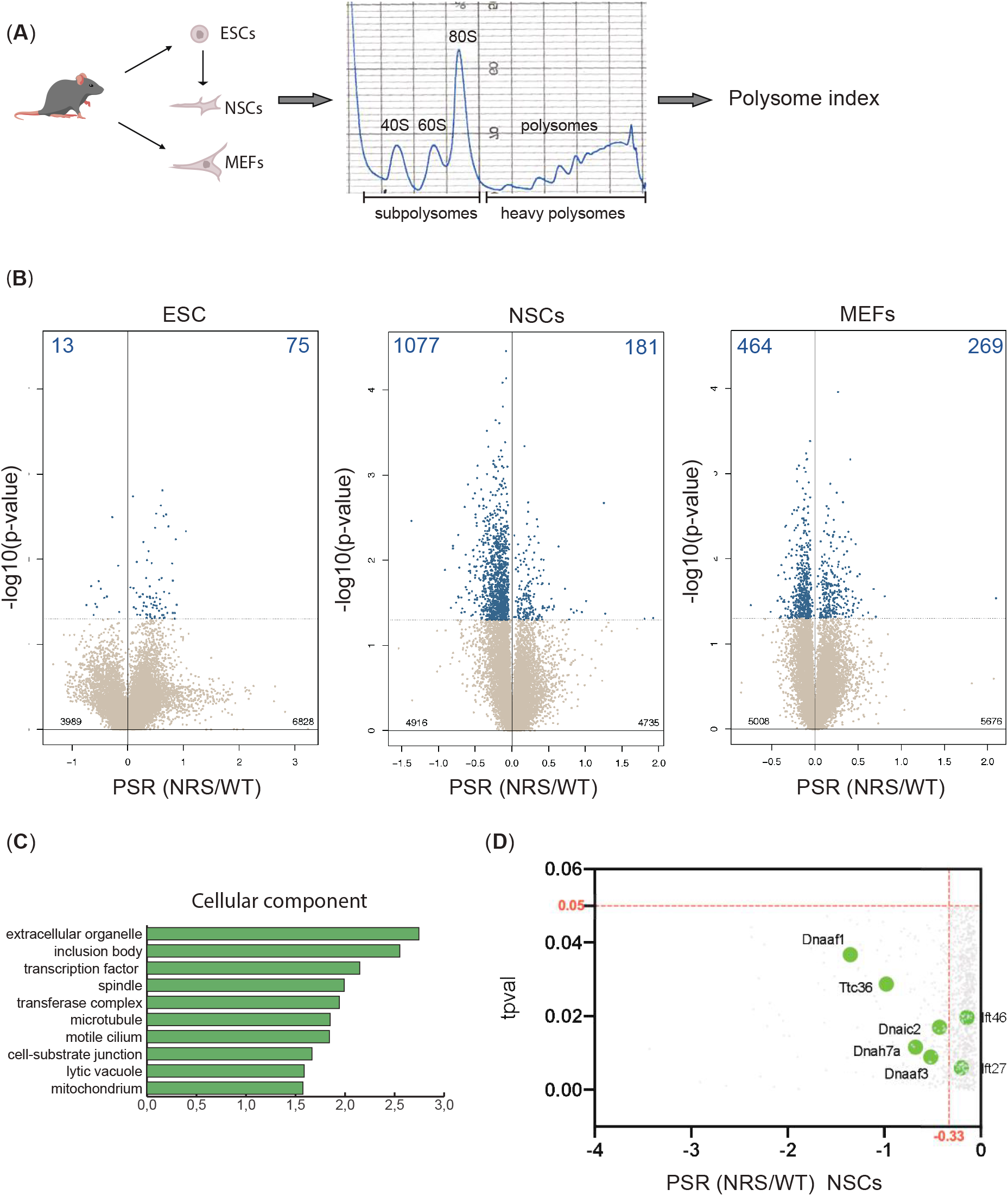
Lack of cytoplasmic SRSF1 results in gross changes in translation. (**A**) Schematic of the experimental approach used to identify translation profiles of wild-type SRSF1 or SRSF1-NRS-expressing ESCs, NSCs and MEF cells. A summary of a fractionation profile is depicted. Absorbance at 254 nm was monitored. RNA isolated from the pooled subpolysomal and polysomal fractions was subjected to RNA sequencing. (**B**) Scatter plots showing the distribution of genes expressed in all cell lines according to their polysome shift ratio (PSR). PSR is calculated as a ratio of polysome index in *Srsf1^+/+^* vs *Srsf1^NRS/NRS^* cells. Blue dots indicate significant changes. Number of significant changes is indicated in the top corners of each plot. (**C**) GO term overrepresentation analysis identifies spindle and motile cilium as enriched cellular components category in the downregulated PSR gene list. (*D*) Plot showing the distribution of genes expressed in NSCs with PSR<0. Selected cilial genes are highlighted.

To confirm that the observed changes are indeed translational and not simply due to gene expression changes, we compared differences in alternative splicing and cytoplasmic RNA expression levels with the observed translational changes. Specifically, we compared genes with significantly different PSR between *Srsf1^+/+^* and *Srsf1^NRS/NRS^* cultures to their respective total mRNA and mRNA isoforms levels from RNA-seq data. Changes in polysome profiling and RNA levels did not correlate in either MEFs or NSCs (***Figure 4-figure supplement 3A***). This indicates that translational changes are a direct response to the loss of SRSF1 in cytoplasm rather than a consequence of altered transcription or mRNA export. Moreover, only a small fraction of differentially expressed splicing isoforms in the cytoplasm (***Figure 4-figure supplement 3B***) were found to differentially translated (***Figure 4-figure supplement 3C).*** Overall, these findings show that the absence of SRSF1 from the cytoplasm has a major impact on translation and that these changes are dependent on the cell-type even though they primarily affect non-cell-type specific RNAs. Altogether, this shows that the SRS1-mediated translation, which was initially observed *in vitro* and cells in culture, does indeed affect a large subset of mRNAs in the mouse. More importantly, it shows that restricting SRSF1 to the nucleus results in perinatal phenotypes, highlighting the need for this activity in a living organism.

Next, we performed gene ontology (GO) analysis to determine whether any functional gene category was differentially translated in SRSF1-NRS expressing cells (***Figure 4C***). We considered only transcripts with a change in PSR of >15%. From this analysis, we identified genes associated with spindle category as translationally downregulated in *Srsf1^NRS/NRS^* NSCs, which agrees with our previous findings in SRSF1-overexpressing 293T cells (Maslon et al., 2014) and with the reduced body size of *Srsf1^NRS/NRS^* animals. Interestingly, the functional category related to motile cilium was also enriched among genes depleted from polysomal fractions of *Srsf1^NRS/NRS^* NSCs (***Figure 4C***). This is consistent with hydrocephalus, immotile sperm and abnormally motile tracheal bundles observed in *Srsf1^NRS/NRS^* animals. In addition, NSCs express a number of mRNAs encoding motile cilia specific proteins and these were downregulated in *Srsf1^NRS/NRS^* NSCs polysome fractions (***Figure 4D***). We observed that dynein axonemal assembly factors *Dnaaf1/Lrrc50* and *Dnaaf3*, which are required for the functional assembly of axonemal dynein motors, as well as motor subunits themselves including those of both inner dynein arms (*i.e.* axonemal dynein heavy chain 7 (*Dnah7a*)) and outer dynein arms (*i.e.* dynein axonemal intermediate chain 2 (*Dnai2*)) were under-represented in the heavy polysome fractions of *Srsf1^NRS/NRS^* cultures. Together, these proteins form the functional macromolecular motors that drive ciliary movement, many of which are found mutated in patients with congenital dysfunction of motile cilia, termed primary ciliary dyskinesia (PCD) (Loges et al., 2009, 2008; Mitchison et al., 2012; Zhang et al., 2002). Importantly, the abundance of these mRNAs was not altered in the cytoplasm of *Srsf1^NRS/NRS^* cultures (***Figure 4-source data 2),*** which was further confirmed by direct RT-PCR for *Dnaaf1* and *Dnaaf3* (***Figure 4, figure supplement 1C***). Altogether, this strongly suggests that SRSF1 participates in the translation of components of the motile cilia and could therefore directly underlie the phenotypes observed in *Srsf1^NRS/NRS^* pups. For these reasons we next focused on a detailed characterization of motile cilia phenotypes in *Srsf1^NRS/NRS^* animals.

### Srsf1^NRS/NRS^ animals have compromised cilia motility

Polysome shifts from *Srsf1^NRS/NRS^* cells indicated that the translation role of SRSF1 is required during the execution of developmental programs. Therefore, we conducted a molecular investigation of motile ciliogenesis, in mouse tracheal epithelial cultures (mTECs) derived from sets of *Srsf1^NRS/NRS^* and *Srsf1^+/+^* littermates. These synchronized primary cultures allow the stepwise study of the *de novo* production of multiciliated cells by exposure to air liquid interphase (ALI) and growth factors (Vladar and Brody, 2013). From progenitors bearing a single non-motile primary cilia to fully differentiated epithelial cells bearing hundreds of motile cilia, these cultures faithfully replicate airway developmental programs (Jain et al., 2010) during the course of a few weeks (***Figure 5A***). These cultures allowed us to observe unique features of motile ciliogenesis including the number of cilia formed by centriole amplification as well as changes in cilia content illustrated by movement parameters (Oltean et al., 2018). Motile cilia move randomly from the onset after the rapid burst of cytoplasmic synthesis, assembly and transport of motile specific components into apically docked cilia primordia, but their waveform changes as cilia mature and movement between ciliary bundles becomes coordinated propelling mucus across the surface of these multiciliated epithelia (***Figure 5A—source data 1)***. The complexity and gradual waveform changes depend on multiple components not limited to ciliary motors, inner and outer dynein arms (IDA, ODA) that power ciliary beating, but also include structural support of motile axonemes (Tektins and Nexin-Dynein Regulatory complexes (N-DRC)) (***Figure 5B***). Mutations in those components are associated with PCD in patients (Loges et al., 2009; Mitchison et al., 2012; Wirschell et al., 2013) and in animal models (Tanaka et al., 2004). High-speed video microscopy and immunofluorescence of mTEC cultures during differentiation indicate that although the production of multiciliated cells is not affected (***Figure 5-figure supplement 1A),*** a significant proportion of cilia remain immotile in *Srsf1^NRS/NRS^* inserts well until ALI13 (***Figure 5C***). Moreover, at these later stages, motile ciliary bundles still present altered beat patterns in relation to the control cultures, as illustrated by kymographs, ciliary beat frequencies (***Figures 5D** and **E***) and beat coordination between ciliary bundles (***Figure 5-figure supplement 1B***). To investigate changes in protein levels during these differentiation stages, we first performed unsupervised analyses of total proteomes from day 4 post-airlift (ALI4) to day 18 (ALI18) using the subset of proteins, which have been annotated with the GO term cilia. We found some cilia candidates that were concurrently and significantly upregulated at later stages of differentiation in *Srsf1^+/+^* controls that failed to be induced in a coordinated manner in *Srsf1^NRS/NRS^* cultures (***Figure 5 F***). Overall, the majority of changes between *Srsf1^NRS/NRS^* and *Srsf1^+/+^* proteomes involved downregulation of proteins with GO terms related to assembly of motile cilia (***Figure 5-figure supplement 1C, D)***. Interestingly, the affected proteins include dynein motor assembly factors (DNAAFs) and axonemal dynein motors (DNAHs), as well as Tektins and N-DRC components that are required for normal axoneme assembly and proper motility of mature ciliary bundles (***Figure 5G***). The abnormal levels of motile cilia components found in *Srsf1^NRS/NRS^* proteomes are consistent with the observed ciliary dysfunctions and structural defects observed by TEM (***Figure 3E*)**.

**Figure 5.**
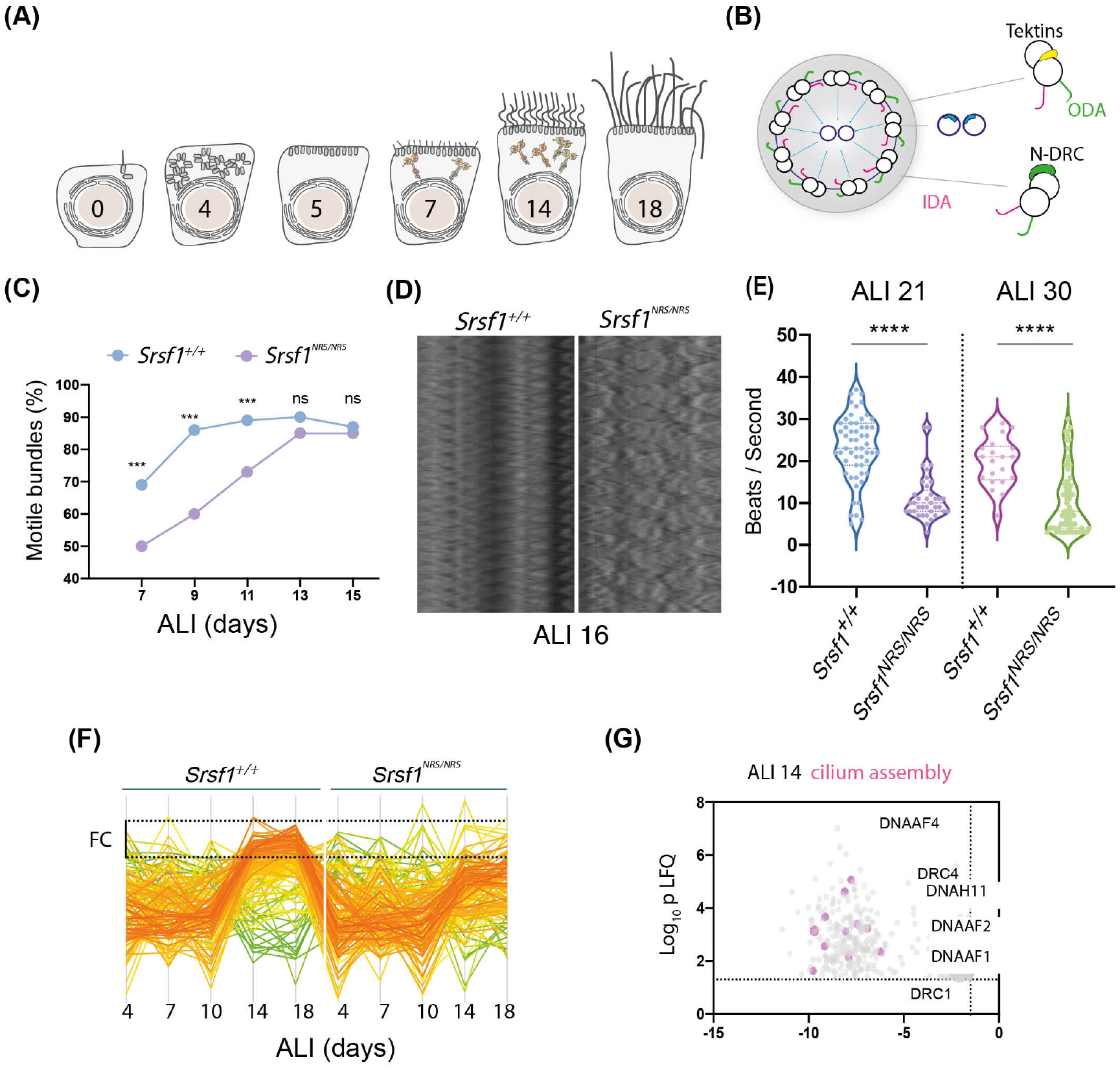
Molecular studies of *Srsf1^NRS/NRS^* mice are consistent with defects in ciliary motility. (**A**) Diagram depicts landmark events in the maturation of motile cilia in mTECs, upon culture in Air Liquid Interface (ALI) as days post airlift. Stages depicted include: centriole amplification and apical docking (ALI 4-6), burst of synthesis and assembly of motile ciliary machinery concomitant with growth and ciliary elongation (ALI 7-14) continuous ciliary beat maturation (ALI 18). (**B**) Diagram of a motile axoneme and some auxiliary components. N-DRC: Nexin-dynein regulatory complexes, ODA: outer dynein arm IDA: inner dynein arm. (**C**) Percentage of motile ciliary bundles during mTEC differentiation. N = 2 animals / genotype at ALI7 and ALI9; N=3 at all later stages. More than >100 ciliary bundles were scored at each time point. Asterisks represent p-value obtained from Chi square exact test at each stage. (**D**) Representative Kymographs of motile bundles illustrate differences in beating amplitude at stages late stages, ALI 16. (**E**) Cilia beat frequency of *Srsf1^NRS/NRS^* and *Srsf1^+/+^* tracheal cultures grown in parallel at at the indicated differentiation stages. Asterisks denote p < 0.001 determined by Mann Whitney tests as *Srsf1^NRS/NRS^* ciliary movement do not follow a normal distribution. (**F**) Z-normalized intensities of proteins containing “cilia” within their GO term aligned along tracheal stages (ALI4-18) and genotypes. Each line represents a single protein, where colour coding denotes those that match closely to the mean trajectory of the group (red) from those that deviate (green). This shows that *Srsf1^+/+^* induces the coordinated expression of multiple cilia-associated proteins during maturation with a greater amplitude that *Srsf1^NRS/NRS^*. Note the tighter distribution of trajectories, and greater fold change (FC) in *Srsf1^+/+^*samples. (**G**) Volcano plot of proteins that are significantly underrepresented in *Srsf1^NRS/NRS^* cultures at ALI14. Pink dots represent proteins with cilia in their GO term.

In summary, we show that the *Srsf1^NRS/NRS^* mouse model mimics organismal and cellular phenotypes attributable to defects in motile cilia, such as those observed in PCD patients. These phenotypic changes are consistent with a lack of SRSF1-mediated cytoplasmic activities that particularly impact on ciliary mRNA, in agreement with altered proteomes and axonemal ultrastructure changes in *Srsf1^NRS/NRS^* animals.

## Discussion

Shuttling SR proteins, including SRSF1, act to co-ordinate nuclear and cytoplasmic steps of gene expression (Botti et al., 2017; Michlewski et al., 2008; Müller-McNicoll et al., 2016). Indeed, SRSF1 promotes translation of mRNAs encoding RNA processing factors and cell-cycle and centrosome-associated proteins (Fu et al., 2013; Maslon et al., 2014; Michlewski et al., 2008). However, despite increasing evidence for post-splicing activities of shuttling SR proteins, their physiological relevance in a living organism has remained enigmatic. Here, we set out to comprehensively investigate the *in vivo* roles and requirement for cytoplasmic SRSF1. We created a mouse model in which SRSF1 is retained in the nucleus, by targeting a nuclear retention signal (NRS) to SRSF1 resulting in a fusion protein (SRSF1-NRS). In our model, abrogation of cytoplasmic SRSF1 function resulted in several deleterious phenotypes without affecting the nuclear roles of SRSF1. To our knowledge, this is the first demonstration that interfering with the nucleo-cytoplasmic shuttling of an RBP has severe phenotypic consequences giving rise to perinatal phenotypes.

### Multiple postnatal phenotypic changes are observed in mice lacking cytoplasmic SRSF1

*Srsf1^NRS/NRS^* mice developed hydrocephalus with variable severity and were smaller than littermate counterparts. Homozygous males also had an increased proportion of immotile sperm, whilst those remaining, exhibit abnormal motility. Moreover, primary cells from *Srsf1^NRS/NRS^* airways and spermatozoa displayed reduced cilia beat frequency, disturbed waveforms and significantly reduced protein levels of motile ciliary components (***Figure 5***). These traits and molecular changes are characteristic of PCD, a genetically heterogenous disease of motile cilia (Lobo et al., 2015). Cilia are small microtubule-based projections found on the surface of most mammalian cells types. Defects in cilia function or structure result in a growing spectrum of diverse human diseases known as the ciliopathies (Reiter and Leroux, 2017). Whilst we observed no gross phenotypes associated with defects in primary (immotile) cilia in *Srsf1^NRS/NRS^* animals, multiple organs that depend on motile cilia were affected. Motile cilia are essential for providing coordinated mechanical force to drive sperm movement or movement of fluids across tissues affecting brain ventricles, trachea and spermatozoa. The constellation of phenotypes, abnormal ciliary ultrastructure, altered proteomes and disrupted motility provide strong evidence that cytoplasmic functions for SRSF1 are essential *in vivo* during motile cilia differentiation.

### Lack of cytoplasmic SRSF1 represses cilia-related mRNA transcripts

We provided evidence showing that abrogation of the cytoplasmic function of SRSF1 led to a decreased polysomal association of a subset of mRNAs in *Srsf1^NRS/NRS^* NSCs, yet its impact on alternative splicing (nuclear function of SRSF1) or on the abundance of target mRNAs in the cytoplasm (mRNA export) was minimal. This strongly suggests that mRNA translation is specifically debilitated when SRSF1 is absent from the cytoplasm. We found that the lack of cytoplasmic SRSF1 had little effect on translation in ESCs, but it affected translation of over one thousand genes in NSCs. These results are consistent with findings that human ESCs maintain low overall levels of translation, with a global increase in high polysomes upon differentiation towards neural lineages (Blair et al., 2017) and thus suggest that SRSF1 participates in developmental programs that rely heavily on translational control.

To further investigate the effects of depletion of cytoplasmic SRSF1 on developmental proteomes, we used tracheal cultures as a model for the stepwise differentiation of motile ciliogenesis that could contribute to the phenotypes observed in *Srsf1^NRS/NRS^* animals. Given the link to SRSF1 dependent translation and mitotic functions at centrosomes in dividing cells (Maslon et al., 2014), it was possible that differentiation of multiciliated cells is impacted, given the hundreds of centrioles destined to become the basal bodies of motile cilia need to be generated within a single postmitotic cell (Meunier and Azimzadeh, 2016). In particular, depletion of mitotic oscillator component CDK1, previously shown to be a direct target of SRSF1 translation (Maslon et al., 2014), hinders differentiation of multiciliated ependymal cells at the stage of massive centriole production (Al Jord et al., 2017). However, in the absence of cytoplasmic SRSF1, differentiation appeared to occur on time and give rise to appropriately long multiciliated bundles (***Figure 5—figure supplement 1).*** However, cilia motility and movement coordination was specifically compromised, and protein levels of motile ciliary components were significantly reduced (***Figure 5F,G***). Furthermore, motility of the sperm flagella is also affected in *Srsf1^NRS/NRS^* animals (***Figure 3C,D***).

During multiciliogenesis, cells transition from bearing a single primary cilium to hundreds of motile cilia requiring *de novo* synthesis of millions of motor subunits and motile ciliary components in the correct stoichiometry within a short developmental window couple to ciliary elongation. Although a well-defined transcriptional code for induction of cilia motility exists, these transcriptional regulators of multiciliogenesis affect hundreds of motile ciliary genes (Lewis and Stracker, 2020) and lead to a drastic decrease in the production of ciliated bundles and cilia number per bundle (Gomperts et al., 2004). These traits were not observed in *Srsf1^NRS/NRS^*cultures. Recent evidence suggests the interesting possibility that spatially regulated translation may be occurring during motile ciliogenesis. During spermatogenesis, *Drosophila* mRNAs encoding motile ciliary proteins are stored in large cytoplasmic granules until their translation is required in the growing sperm axoneme, possibly regulating their translation (Fingerhut and Yamashita, 2020). In vertebrates, there is evidence of phase-separated cytoplasmic foci termed dynein axonemal particles (DynAPs) that contain assembly factors, chaperones and axonemal dynein proteins, that may provide a favorable environment for their orderly assembly and final transport onto cilia (Huizar et al., 2018). It remains to be seen whether this is also the case for DynAPs, but as with most other phase-separated condensates, RNA is key to regulate the formation, size and composition of these phase transitions (Langdon et al., 2018; Maharana et al., 2018). It is possible that SRSF1 could contribute to localized mRNA delivery and/or translation at these cytoplasmic foci. It could promote the formation of mRNPs that confine such regulated transcripts until a timely co-translational assembly of these large protein complexes is required during motile ciliogenesis.

An intriguing possibility is that SRSF1 is able to respond to metabolic demands of the cell needing to reshape its proteome during differentiation. We propose that cytoplasmic activities of SRSF1 are important for the execution of cilia motility programs, providing an extra regulatory level that will ensure responsive and scaleable translation at the peak demand of assembly of the complex motile cilia machinery.

In summary, we have developed the first mouse model that allowed us to assess the *in vivo* relevance of SRSF1 nucleo-cytoplasmic shuttling (***Figure 6***). We conclude that the presence of the splicing factor SRSF1 in the cytoplasm is essential for proper postnatal development, and this is mostly due to the effect of SRSF1 in mRNA translation, which critically affects developmental programs and target mRNAs involved in motile cilia biogenesis and function.

**Figure 6.**
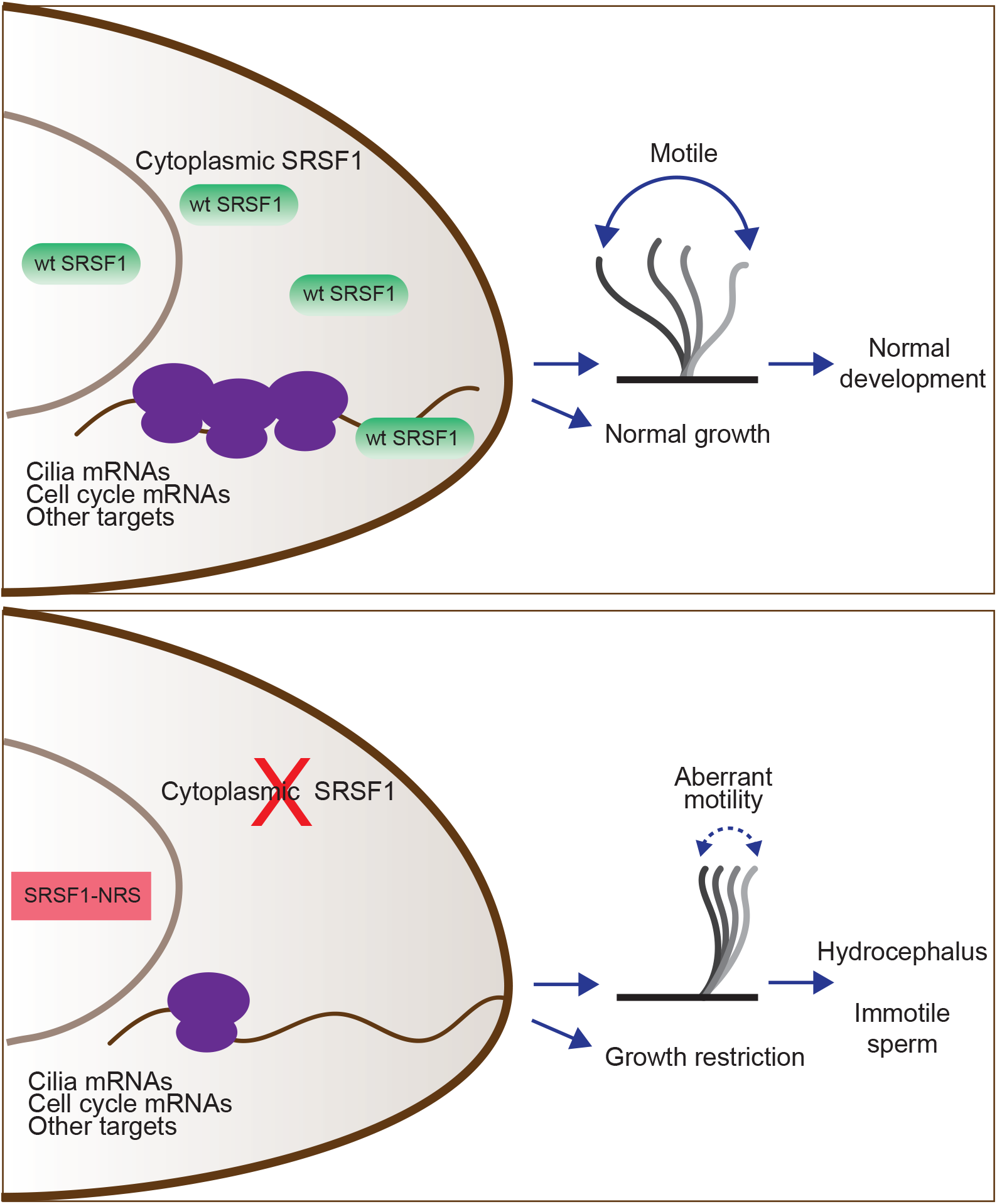
Restricting SRSF1 to the nucleus results in perinatal phenotypes in the mouse in particular affecting motile cilia. This highlights the physiological role of splicing factor nucleo-cytoplasmic shuttling to reprogram gene expression networks to meet high cellular demands.

## Material and methods

### Key resources table

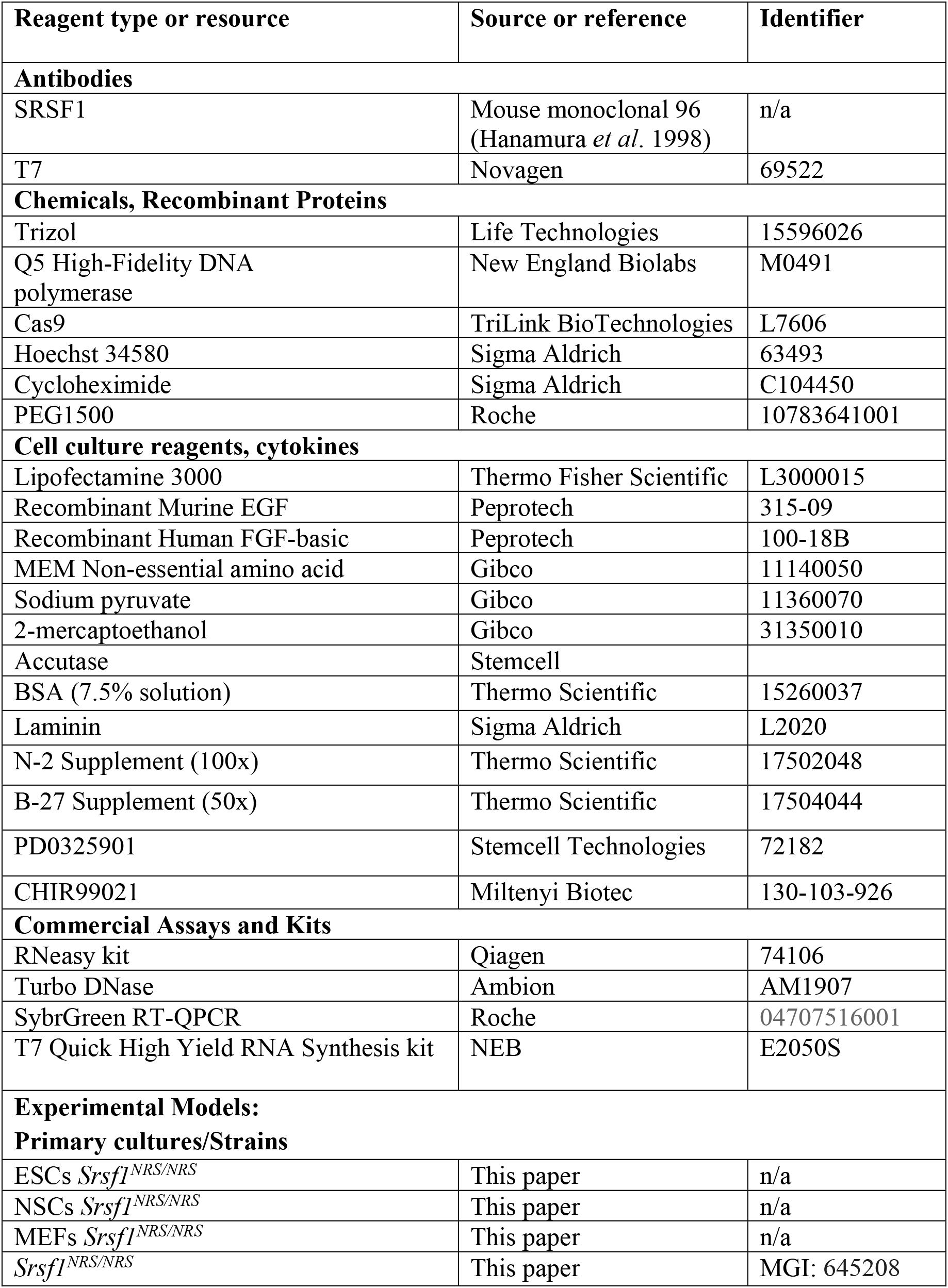
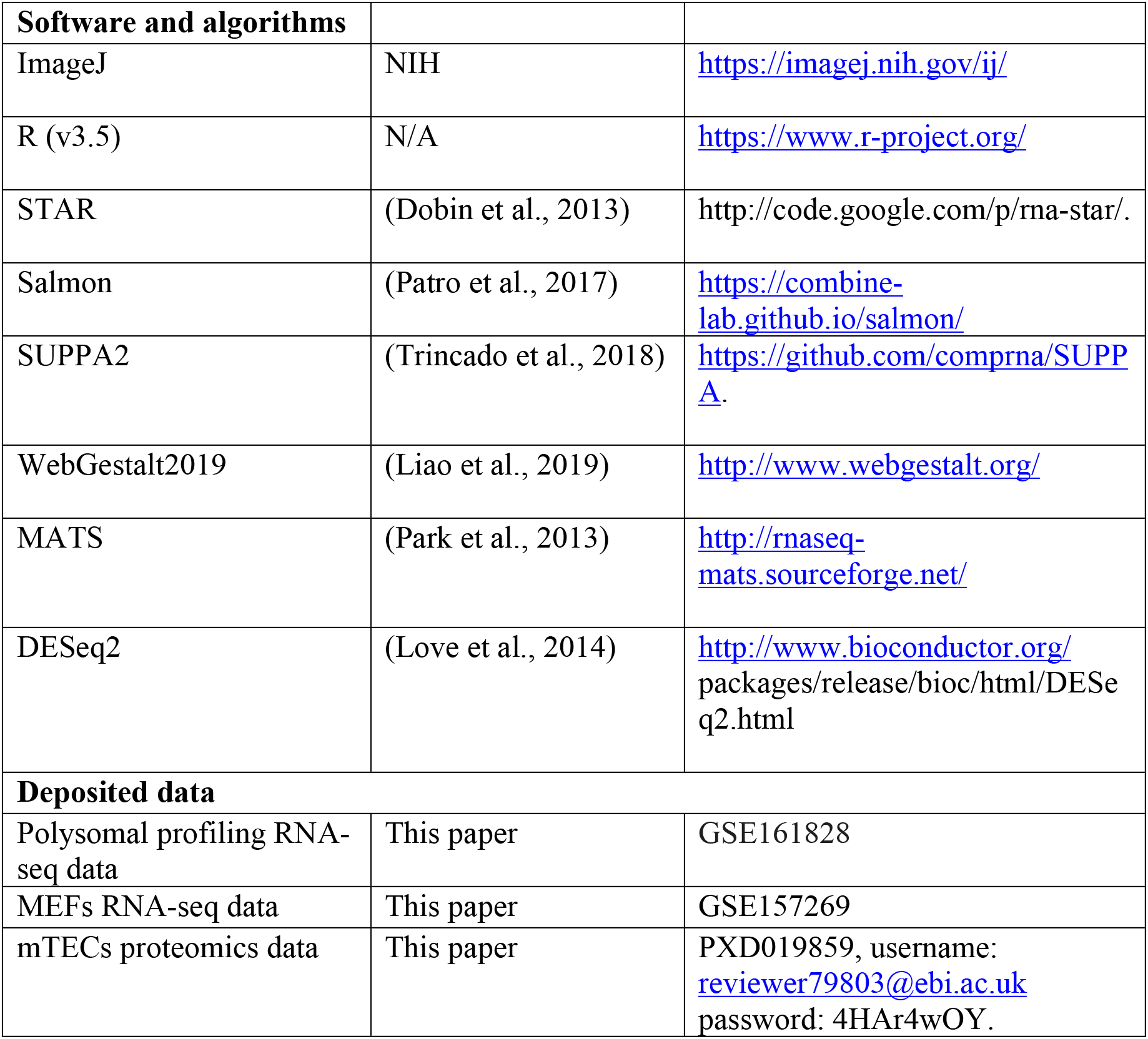

### Animal experiments

We followed international, national and institutional guidelines for the care and use of animals. Animal experiments were carried out under UK Home Office Project Licenses PPL 60/4424, PB0DC8431 and P18921CDE in facilities at the University of Edinburgh (PEL 60/2605) and were approved by the University of Edinburgh animal welfare and ethical review body.

### CRISPR/Cas9 Gene Editing in Mouse Zygotes

The CRISPR target sequence (20-nucleotide sequence followed by a protospacer adjacent motif (PAM) of ‘NGG’) were selected using the prediction software (http://www.broadinstitute.org/rnai/public/analysis-tools/sgrna-design). Single gRNAs targeting exon 4 of *Srsf1* (Srsf1-205, ENSMUST00000139129.8) were annealed and cloned into the PX458 plasmid (Cong et al., 2013). The guide region was then amplified by PCR and paired guide RNAs synthesised by in vitro transcription (T7 Quick High Yield RNA Synthesis kit, NEB, #E2050S). Single stranded DNA oligonucleotides (WT oligo: “CGTAGCAGAAGCAACAGCAGGAGTCGCAGTTACTCCCCAAGGAGAAGCAGAGGATCACCACGCTATTCTCCCCGTCATAGCAGATCTCGCTCTCGTACATAAGATGATTGATGACACTTTTTGTAGAACCCATGTTGTATACAGTTTTCCTTTACTCAGTACAATCTTTTCATTTTTTTAATTCAAGCTGTTTTGTTCAG”, NRS oligo: “AAGCAGAGGATCACCACGCTATTCTCCCCGTCATAGCAGATCTCGCTCTCGTACAGGATCCCCTCCGCCCGTGTCGAAGCGAGAGTCCAAGTCTAGGTCGCGGTCCAAGAGCCCACCCAAGTCTCCAGAAGAAGAGGGAGCAGTTTCTTCCATGGCATCGATGACAGGTGGCCAACAGATGGGTTAAGATGATTGGTGACACTTTTTGTAGAACCCATGTTGTATACAGTTTTCCTTTACTC” and T7 oligo: “TTACTCCCCAAGGAGAAGCAGAGGATCACCACGCTATTCTCCCCGTCATA GCAGATCTCGCTCTCGTACAGGAT**CCCCCGGCGCCGGCG**CCATGGCATC G ATGACAGGTGGCCAACAGATGGGTTAAGATGATTGGTGACACTTTTTGTAGAACCCATGTTGTATACAGTTTTCCTTTACTCAGTACAATCTTTTCA”) were synthesized by IDT. The sequence encoding a small peptide linker (P-G-A-G-A), inserted between the NRS and the T7 peptide, is highlighted in bold. Gene editing was performed by microinjection of RNA encoding the wild-type Cas9 nuclease (50 ng/μl, TriLink BioTechnologies, #L7606), 25 ng/μl single guide RNA (sgRNA) targeting exon

4 of *Srsf1*, and 150 ng/μl of both NRS-T7 and WT single-stranded DNA oligonucleotides of a similar length containing 50-100 nucleotides of homology flanking both sides of the sgRNA as repair templates into (C57BL/6 x CBA) F2 zygotes (Crichton et al., 2017). The injected zygotes were cultured overnight in KSOM for subsequent transfer to the oviduct of pseudopregnant recipient females (Joyner, 2000). From microinjection CRISPR targeting, 57 pups were born, and genomic DNA from ear-clip tissue was subject to PCR screening and subsequent sequence verification of successful mutagenesis events by Sanger sequencing ***(Figure 1—figure supplement 1B***). Two heterozygous founder mice were each backcrossed for one generation to C57BL/6 wild type mice to allow unwanted allelic variants to segregate. Of the resulting 32 pups, 10 animals, were confirmed by sequencing to be heterozygous for the correct SRSF1-NRS allele (***Figure 1—figure supplement 1B***, Allele Accession ID: MGI:6452080) and subsequently intercrossed. Genotyping was performed by PCR using the following forward and reverse primers: TTGATGGGCCCAGAAGTCC and ATAGGGCCCTCTAGACAATTTCATCTGTGACAATAGC, respectively.

### Srsf1^NRS^ mice

Two heterozygous *Srsf1^NRS^* founder mice were each backcrossed for one generation to C57BL/6 wild type mice to allow unwanted allelic variants to segregate. Of the resulting 32 pups, 10 animals, were confirmed by sequencing to be heterozygous for the correct SRSF1-NRS allele (***Figure 1#x2014;figure supplement 1B***, Allele Accession ID: MGI:6452080) and subsequently intercrossed. The phenotypes reported in Figs. 1-3 were present in *Srsf1^NRS/NRS^* animals derived from both independent founder mice, confirming their association with homozygosity for *SRSF1^NRS^* rather than an off-target or spontaneous mutation. For subsequent studies *SRSF1^NRS^* heterozygous animals were crossed onto the ubiquitously expressing ARL13B-Cerulean biosensor (Ford et al., 2018). In this background *Srsf1^NRS/NRS^* mutants were born at the frequencies of 32 *Srsf1^+/+^*, 75 *Srsf1^+/NRS^*, 27 *Srsf1^NRS/NRS^*. Initial genotyping was performed by PCR using the following forward and reverse primers: TTGATGGGCCCAGAAGTCC and ATAGGGCCCTCTAGACAATTTCATCTGTGACAATAGC, respectively. The genotyping of the compound animals was done by TransNetyx.

### Histology

Whole brains from P14 mice were embedded in paraffin wax and 5μM sections cut using a microtome before oven drying overnight at 60°;C. Sections were dewaxed in xylene and rehydrated in decreasing concentrations of ethanol with a final rinse in running water. Sections were stained in haematoxylin for 4 min, rinsed in water then placed in 1% HCl in 70% EtOH for 5 sec before rinsing again in water and treating with Lithium carbonate solution for 5 sec. Tissue sections were rinsed well in running water for 5 min and stained in Eosin for 2 min before rinsing in water and washing three times in 100% EtOH for 1 min each. Sections were cleared in fresh xylene 3 times for 5 min and mounted using DPX mounting media.

### Heterokaryon assay

This was performed as previously described (Cazalla et al., 2002; Piñol-Roma and Dreyfuss, 1992), with minor modifications. Donor mouse NSCs were seeded on laminin-coated cover slips, followed by co-incubation with an excess of HeLa cells (recipient) for 3 h in the presence of 50 μg/ml cycloheximide (Sigma, # C104450), to prevent further protein synthesis in the heterokaryons. The concentration of cycloheximide was then increased to 100 μg/ml, and the cells were incubated for an additional 30 min prior to fusion. Cells were fused with PEG1500 (Roche, #10783641001) in the presence of 100 μg/ml cycloheximide and the heterokaryons were further incubated for 3 h in media containing 100 μg/ml. Cells were then fixed and stained with a T7 antibody, and DNA was stained with Hoechst 33258 (Sigma, #63493) to distinguish between mouse and human nuclei.

### Mouse tracheal epithelial cultures

These were established as described previously (Vladar and Brody, 2013). *Srsf1^+/+^* and *Srsf1^NRS/NRS^* animals were always sacrificed as pairs. Tracheas were dissected and dissociated individually. More than 5 pairs of animals were used to establish tracheal cultures used along this study. Cells were expanded into monolayers of progenitor cells in T25 flasks until confluency and dissociated as described to be seeded onto 6-9 transwells of 6.5 mm each depending on cell density. Under these conditions, our cultures have roughly no ciliary bundles until day 4 and visibly motile cilia until day 7, reaching mature coordinated beating between adjacent bundles by 25-30 days post-airlifting. Membranes of transwells were released from inserts and inverted onto a glass bottom plate for imaging after which cells were harvested for proteomics, immunofluorescence or electron microscopy.

### Protein extraction, antibodies and Western Blotting

Cell pellets were lysed in 50 mM Tris pH8.0, 150 mM NaCl, 1 % NP-40 buffer containing protease inhibitors. Protein samples either from ESCs or liver extracts were separated by SDS-PAGE and electroblotted onto nitrocellulose membranes (Whatman) using iblot System for 6 min (Invitrogen). Non-specific binding sites were blocked by incubation of the membrane with 5 % nonfat milk in PBS containing 0.1% Tween 20 (PBST). Proteins were detected using the following primary antibodies diluted in blocking solution: mouse monoclonal anti-SRSF1 (clone 96; 1:1000 (Hanamura et al., 1998)), mouse monoclonal anti-tubulin (Sigma #T8328)) and mouse monoclonal anti-T7 (Novagen, #69522, 1:10,000). Following washing in PBST, blots were incubated with the appropriate secondary antibodies conjugated to horse-radish peroxidase (Pierce) and detected with Super Signal West Pico detection reagent (Pierce).

### Proteomics

Each transwell (1 experimental replicate) of mTECs cultures were lysed in 350 μl of PBS, 2% SDS and antiproteolytic cocktail, kept at −80°C until all time points were harvested to be run simultaneously. Total proteomes were derived from two animals/genotype with 3 experimental replicate per time point (Day 4 to 10, animal pair 1; day 14 to 18 animal pair two). Cell lysates were digested using sequential digestion of LysC (Wako) and trypsin (Pierce) using the FASP protocol (https://pubmed.ncbi.nlm.nih.gov/25063446/). Samples were acidified to 1% TFA final volume and clarified by spinning on a benchtop centrifuge (15k g, 5 min). Sample clean-up to remove salts was performed using C18 stage-tips (Rappsilber et al., 2003). Samples were eluted in 25 μl of 80% Acetonitrile containing 0.1% TFA and dried using a SpeedVac system at 30°C and resuspended in 0.1% (v/v) TFA such that each sample contained 0.2 μg/ml. All samples were run on an Orbitrap FusionTM Lumos mass spectrometer coupled to an Ultimate 3000, RSL-Nano uHPLC (both Thermo Fisher). 5 ul of the samples were injected onto an Aurora column (25 cm, 75um ID Ionoptiks, Australia) and heated to 50C. Peptides were separated by a 150 min gradient from 5-40% Acetonitrile in 0.5% Acetic acid. Data were acquired as data-dependent acquisition with the following settings: MS resolution 240k, cycle time 1 s, MS/MS HCD ion-trap rapid acquisition, injection time 28ms. The data were analyzed using the MaxQuant 1.6 software suite (https://www.maxquant.org/) by searching against the murine Uniprot database with the standard settings enabling LFQ determination and matching. The data were further analyzed using the Perseus software suite. LFQ values were normalized, 0-values were imputed using a normal distribution using the standard settings. To generate expression profiles of cilium-related proteins, gene ontology annotations were added and all entries containing “cilia” were retained. Expression profiles were normalized (using Z-score) and expression profiles were plotted.

### Cilia motility analysis

mTECs were imaged by releasing the membrane from each transwell on which they were grown and inverting it onto a glass bottom plate. Ciliated bundles were imaged randomly as they appeared on the field of view using a 60x Plan Apochromat VC 1.2 WI DIC N2 lens within a preheated chamber at 37 °C. Motile vs immotile bundles were scored by direct visual inspection of these movies. The number of all counts for each category at each time point were used to calculate the significance of the different proportions between *Srsf1^NRS/NRS^* and *Srsf1^+/+^* cultures by Fisher Exact test. Other parameters of cilia motility were analyzed in FIJI as indicated in each panel by slowing down animation to manually count beats/sec of distinct ciliary bundles or automatically by the custom-written ImageJ plugin “Cilility_JNH” (available upon request). The underlying analysis method was adapted from (Olstad et al., 2019) and is based on the periodic changes of pixel intensities in the image caused by ciliary beating that are used by the plugin to determine ciliary beat frequency. The intensity time course at each pixel in the image was converted into a frequency spectrum using a Fast-Fourier-Transformation (code: edu.emory.mathcs.jtransforms.fft by Piotr Wendykier, Emory University). Each spectrum was smoothed with a user-defined averaging sliding window (5 Hz). The position and power of the highest peak in the spectrum within a user-defined range was determined (will be referred as primary frequency). The user-defined range was defined as 3 to 62.5 Hz, which represents a quarter of the acquisition frequency (250 Hz). Next, a custom noise-threshold algorithm was applied to separate the image into noise and signal regions: (1) Pixels were sorted by the power of the primary frequency; (2) The average and standard deviation (SD) of the 20% of pixel with the lowest power were determined and used to define a power threshold as average + 1.5x SD; (3) all pixels with a primary frequency power above the threshold were considered as signal pixels, all other pixels were considered as noise pixels. A “noise” power spectrum was determined as the average + 1.5x SD of all power spectra from pixels belonging to noise regions. The “noise” power-spectrum was subtracted from the power spectrum at all “signal” pixel positions (power below zero was set to zero). For each cell, the resulting power spectrum from all “signal” pixels was averaged in to a “signal” power spectrum. The frequency of the highest peak in the “signal” power spectrum within the user-defined range of 3 to 62.5 Hz determined the ciliary beat frequency. All results were scrutinized by a trained observer.

### Color-coded time projections of swimming mouse sperm

To generate color-coded time projections of time-lapse images of mouse sperm, images were processed as follows in FIJI (Schindelin et al., 2012). They were pre-processed using SpermQ_Preparator (Hansen et al., 2018): (i) pixel intensities were inverted; (ii) pixel intensities were rescaled so that intensities in the image covered the whole bit range; (iii) the image stack was blurred with a Gaussian blur (sigma = 0.5 px) to reduce noise; (iv) a median intensity projection of the image stack was subtracted from all images in the stack to remove the static image background; (5) pixel intensities were again rescaled so that intensities in the image covered the whole bit range; (6) the background was subtracted using ImageJ’s ‘Subtract Background’ method (radius 5 px). Next, a selected time span (time indicated in the figure legend) was converted into a color-coded time projection using FIJI’s function ‘Temporal-Color Code’ (Schindelin et al., 2012).

### Cell cultures

ESCs were grown on gelatin-coated plates in 2i media: 1:1 Neurobasal and DMEM/F12, supplemented with 0.5X N2 (Thermo Scientific, #17502048), 0.5x B27 (Thermo Scientific, #17504044), 0.05% BSA (Thermo Scientific, #15260037), 1mM PD0325901 (Stemcell Technologies, #72182), 3μM CHIR99021 (Miltenyi Biotec, #130-103-926), 2mM L-glutamine, 0.15mM monothioglycerol, 100U/ml LIF). NSCs were grown on laminin-coated (Sigma Aldrich, #L2020) plates in DMEM/F12 medium supplemented with 2mM L-glutamine, 0.5x N2, B27, glucose, BSA, HEPES and 10ng/ml of both mouse EGF (Peprotech, #315-09) and human FGF-2 (Peprotech, #100-18B). MEFs were cultured in Dulbecco’s Modified Eagle’s Medium (DMEM) supplemented with 10% fetal bovine serum (FBS).

### Derivation and maintenance of ESCs

Embryonic stem cells were established from *Srsf1^NRS/+^* x *Srsf1^NRS/+^* crosses, following an adapted version of the protocol by (Czechanski et al., 2014). E3.5 blastocysts were isolated and plated in 4-well plates pre-seeded with Mitomycin-C-treated MEFs. All derivation and downstream propagation of established lines was carried out in 2i media. After the first 48 h during which the blastocysts were undisturbed, media was changed every other day. Outgrowths were isolated after approximately one week and transferred to a fresh feeder-coated plate. Successfully derived cells were slowly weaned from feeders by continual passaging before genotyping to avoid contamination of genomic DNA.

### Neuroectodermal specification

ESCs were cultured under feeder-free conditions in 2i medium. One day prior to induction of differentiation cells were seeded at high density in 2i medium. The following day, cells were detached using Accutase (Stemcell), resuspended in N2B27 media (1:1 Neurobasal and DMEM/F12, supplemented with 0.5X N2, 0.5x B27, 0.1 mM 2-mercaptoethanol, 0.2 mM L-glutamine), counted and plated at approximately 10,000 cells per cm^2^ onto either 15 cm plates or 6 well plates that have been coated with a 0.1% gelatin solution. Culture medium was changed every second day.

### Deriving NS cells

For derivation of neural stem cells at day 7 of differentiation, cultures were detached using Accutase, 2-3 x 10^6^ cells were re-plated into an uncoated T75 flask in NS expansion media, comprising DMEM/F12 medium supplemented with 2mM L-glutamine, 0.5x N2, B27, glucose, BSA, HEPES and 10ng/ml of both mouse EGF and human FGF-2. Within 2-3 days, thousands of cell aggregates formed in suspension culture and were harvested by centrifugation at 700 rpm for 1 min. They were then re-plated onto a laminin coated T75 flask. After few days, cell aggregates attached to the flask and outgrew with NS cell.

### RNA isolation and RT-qPCR

RNA was isolated using TRIzol (Life Technologies, #15596026) or RNAeasy (Qiagen, #74106) following the manufacturer’s protocol. RNA was then treated with Turbo Dnase (Ambion, #AM1907) and transcribed to cDNA using First-Strand Synthesis System from Roche. This was followed by SybrGreen detection system (Lightcycler 2x SybrGreen Mix, Roche, #04707516001).

### RNA-Seq analysis

RNA was extracted from *Srsf1^+/+^, Srsf1^NRS/+^* or *Srsf1^NRS/NRS^* MEFs and purified using RNeasy kit from three independent experiments. RNA-seq libraries were generated from Poly(A)^+^ mRNA using TrueSeq protocol and sequenced using the Illumina Hi-Seq 4000 machine (WTCRF Edinburgh) to generate 75 bases, paired-end reads. Reads were mapped to the mouse (mm10) genome. Paired reads were pseudoaligned to the GRCm38 (mm10) Ensembl 87 transcriptome using salmon (Patro et al., 2017). Splicing changes were inferred from transcript TPMs using SUPPA2 (Trincado et al., 2018) from gene definitions in the Ensembl 87. SUPPA2 infers splicing changes (dPSI) from changes in transcript models across the two conditions being compared. These results were filtered on mean transcript expression (TPM>0.5).

### Cell fractionation and sucrose gradient centrifugation

ESCs, MEFs, or NSCs were treated with 50 μg/ml cycloheximide for 30 min. Cells were subsequently washed twice in ice-cold PBS containing cycloheximide. Cytoplasmic extracts were prepared as previously described (Sanford et al., 2004). Sucrose gradients (10-45%) containing 20mM Tris, pH 7.5, 5mM MgCl2, 100mM KCl were made using the BioComp gradient master. Extracts were loaded onto the gradient and centrifuged for 2.5 h at 41,000 rpm in Sorvall centrifuge with SW41Ti rotor. Following centrifugation, gradients were fractionated using a BioComp gradient station model 153 (BioComp Instruments, Inc. New Brunswick, Canada) measuring cytosolic RNA at 254 nm. Fractions eight to eleven (polysomal fractions) and one to seven (subpolysomal fraction) were pooled and diluted sucrose concentration adjusted to 20%. The RNA extraction was performed as described above.

### Polysomal shift analysis

Experiments were performed in three biological replicates. Monosomal, polysomal and cytoplasmic reads were mapped to the mouse genome sequence (mm10) by STAR software (Dobin et al., 2013) (v2.0.7f) with –outFilterScoreMinOverLread 0.3 –out FilterMatchNminOverLread 0.3settings and Ensembl release M21 transcript annotation. Polysomal Indexes (PIs) were calculated for each condition for the set of protein coding genes based on a procedure previously described (Maslon et al., 2014). The transcript abundance metric, RPKM, was replaced by transcripts per million reads (TPMs) as such normalization allows comparison between samples without biases. The PIs and associated polysomal shift ratios (PSRs) were calculated for each sample by pooling reads from replicates. The statistical significance of PSRs were assessed by Student’s t-Test on the variability of PIs of individual replicates. Alternative splicing events were retrieved using rMATS (v3.2.5) (Park et al., 2013). The results were filtered by a minimum coverage of 10 reads per junction and replicate, dPSI>0.2 and FDR<0.05. Differential expression analysis was performed with DEseq2 (v1.26.0) (Love et al., 2014) and results were filtered by FDR<0.05.

### Gene Ontology analysis

GO term enrichment analysis was performed using WebGestalt2019 (Liao et al., 2019). Genes downregulated at least 0.85-fold (PSR<=-0.23) or upregulated at least 1.15-fold were used as an input. The following parameters were used for the enrichment analysis: minimum number of IDs in the category: 5 and maximum number of IDs in the category: 300

### Quantification and Statistical analysis

All statistical analysis was carried out using GraphPad Prism 8 (version 8.4.1; GraphPad software, USA) as described in the text. To determine statistical significance, unpaired *t*-tests were used to compare between two groups, unless otherwise indicated. The mean ± the standard error of the mean (SEM) is reported in the corresponding figures as indicated. Statistical significance was set at *P*<0.05. Fisher’s exact test was used to determine the significance in animal studies and to classify ciliary bundles in motile or immotile during mTEC maturation. All *in vitro* experiments were repeated in three biological replicates and several litters were used for *in vivo* studies, as indicated in each section.

## Supporting information

Supplemental Table 1, related to Figure 4

Supplementary Table 2, related to Figure 4

Supplemental Table 3, related to Figure 4

Supplemental movie 1, related to Figure 5

## Data availability

The accession number for total RNA-seq data related to splicing analysis at Gene Expression Omnibus (GEO) is GSE157269G. Polysomal, monosomal and cytoplasmic RNA-sequencing data are available at GEO with the accession number GSE161828. The mass spectrometry proteomics data are available at ProteomeXchange Consortium via the PRIDE partner repository with the dataset identifier PXD019859. Username; Password:4HAr4wOY. All custom MATLAB scripts and ImageJ plugins are available upon request.

## Acknowledgements

We thank Andrew Wood (MRC HGU) for discussion on CRISPR/Cas strategies, Lisa McKie for Histology work, Jimi Wills for mass spectrometry sample preparation and analysis and the MRC Advanced Imaging Resource for expert advice.

## Author contributions

F.H., M.M.M, I.R.A. and J.F.C. conceived, designed, and interpreted the experiments. F.H., M.M.M. and P.L.Y. performed most of the experiments and data analysis. S.A. and N.B. provided all the bioinformatics analysis and statistical analysis. P.L.Y. and P.M. analyzed all the cilia phenotypes and contributed to discussion and interpretation of the results. J.N.H. and D.W. provided image analysis. A.vK. carried out mass spectrometry data and bioinformatic analysis. Edinburgh University Bioresearch & Veterinary Services performed CRISPR injections and mouse work. The manuscript was co-written by all authors.

## Funding

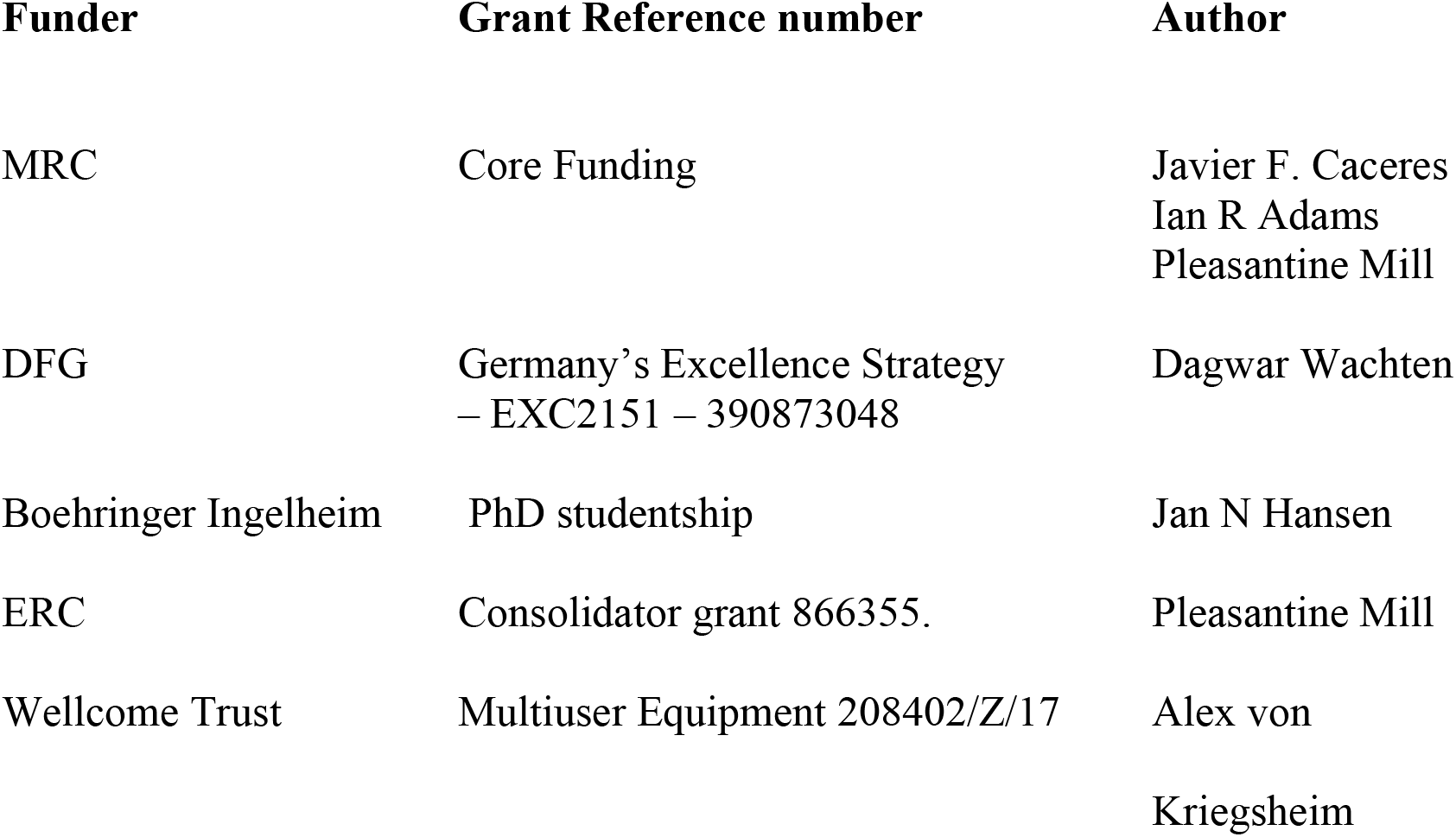

## Figure supplements, related to main Figure

**Figure 1—figure supplement 1**. Genotyping of *Srsf1^NRS/NRS^* mice.

**Figure 1—figure supplement 2**. Subcellular localization of SRSF1 and SRSF1-NRS proteins.

**Figure 3—figure supplement 1**. Live images of ARL13B-Cerulean cilia of ependymal cells

**Figure 4—figure supplement 1**. The lack of cytoplasmic SRSF1 does not affect pre-mRNA splicing or mRNA export.

**Figure 4—figure supplement 2**. Lack of cytoplasmic SRSF1 induces translational changes not restricted to lineage specific transcript.

**Figure 4—figure supplement 3**. PSR values are independent of alternative splicing and gene expression changes in cytoplasm.

**Figure 4—data source 1, Table 1.** Splicing changes between MEF lines derived from *Srsf1^NRS/NRS^* and *Srsf1^+/+^* littermates.

**Figure 4—data source 2, Table 2.** RNA sequencing analysis on cytoplasmic fractions from *Srsf1^+/+^* or *Srsf1^NRS/NRS^* MEFs and NSCs.

**Figure 4—data source 3, Table 3.** PSR values for genes identified in *Srsf1^+/+^* or *Srsf1^NRS/NRS^* ESCs, MEFs and NSCs, respectively.

**Figure 5—figure supplement 1.** Total proteomes of mouse (mTECs)

**Figure 5—data source 1, Movie 1.** Composite image illustrating how ciliary movement of cilia within and between bundles changes as mTECs mature in ALI cultures.

**Figure 1-figure supplement 1.**
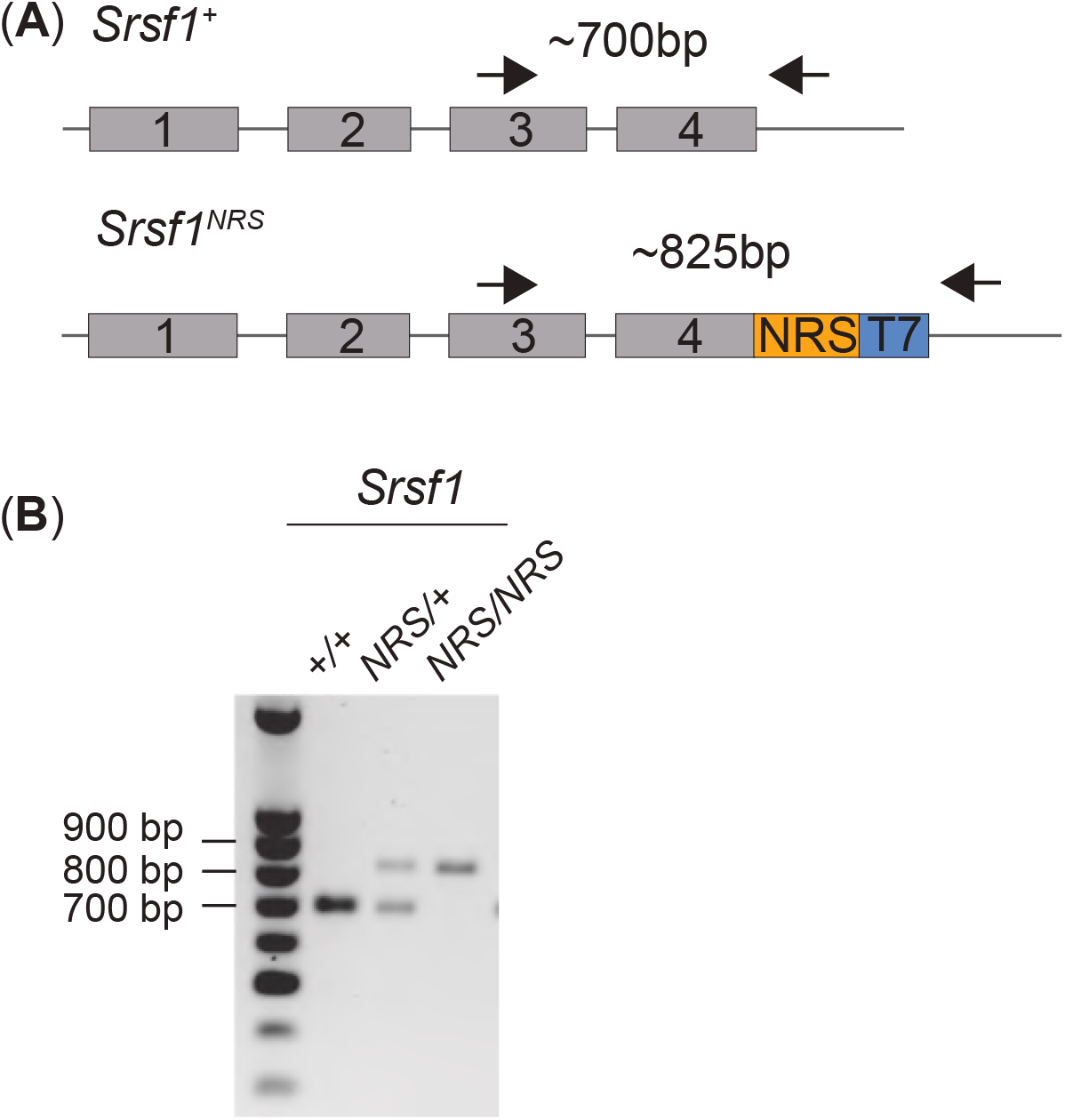
Genotyping of *Srsf1^NRS/NRS^* mice. (**A**) Schematic of the PCR-based genotyping strategy adopted to determine the knock-in status of the C-terminal region of endogenous SRSF1. The arrows represent primers used for PCR. Amplicon sizes for the *Srsf1^NRS^* or *Srsf1^wt^* are shown. (**B**) PCR analysis of genomic DNA of the first two cohorts of pups from heterozygous intercrosses. The PCR products of animals selected for use were all subject to Sanger-sequencing to confirm the presence of an intact *Srsf1^NRS^* allele at the correct genomic locus. Representative samples are shown.

**Figure 1-figure supplement 2.**
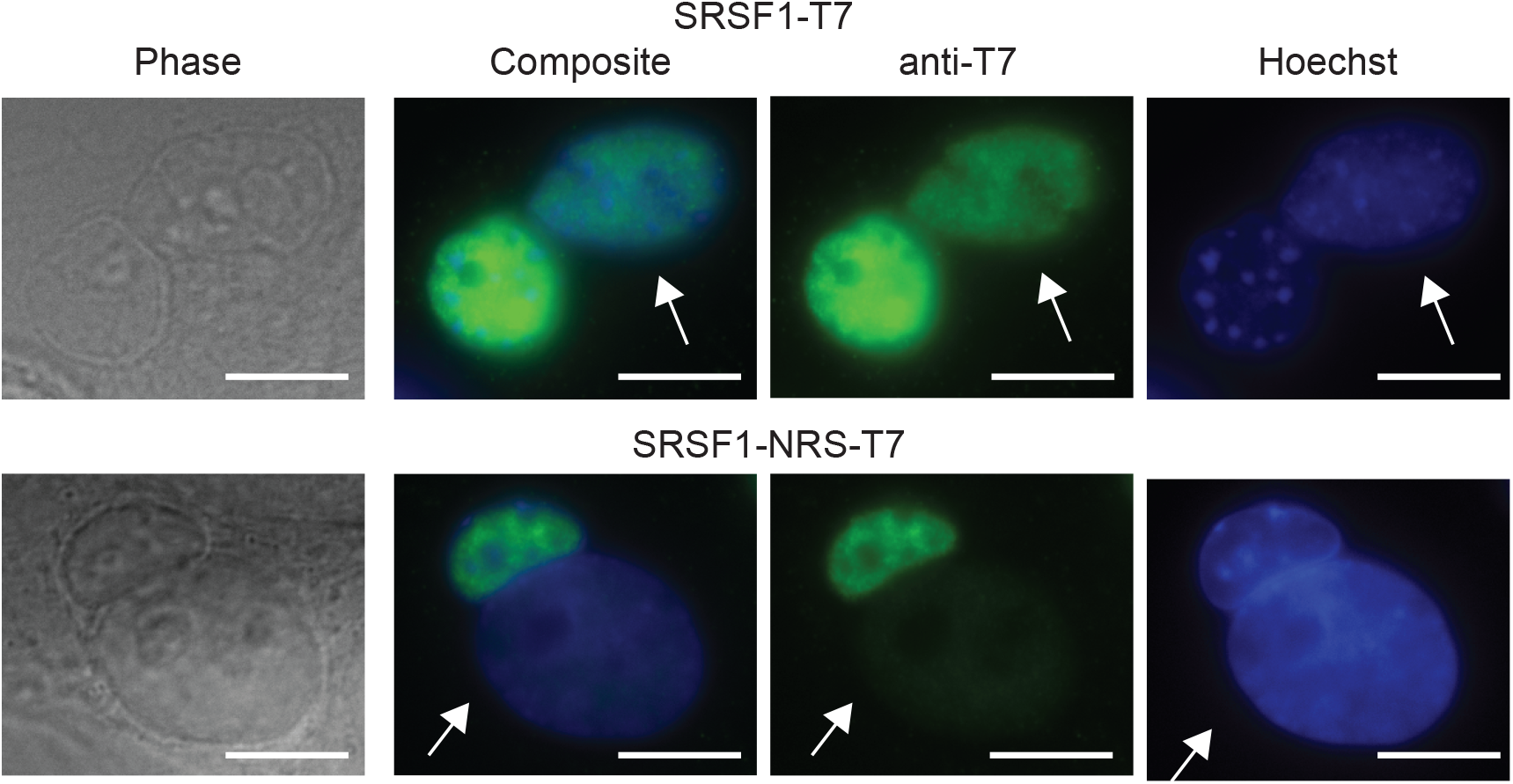
Localization of SRSF1 and SRSF1-NRS protein. Analysis of nucleocytoplasmic shuttling of SRSF1-NRS in NSCs using a heterokaryon assay. Mouse NSCs (donor) transfected with wild type T7-tagged SRSF1 were used as a control (upper panel). Human HeLa cells (recipient) were fused with mouse NSCs (donor) in the presence of cycloheximide to form heterokaryons. The cells were incubated further for 3 h in the presence of cycloheximide and fixed. The localization of SRSF1 was determined with anti-T7 monoclonal antibody. Hoechst 33258 was used to differentially stain mouse and human nuclei. Human (recipient) nuclei are depicted (white arrow). Scale bar, 10μm.

**Figure 3-figure supplement 1.**
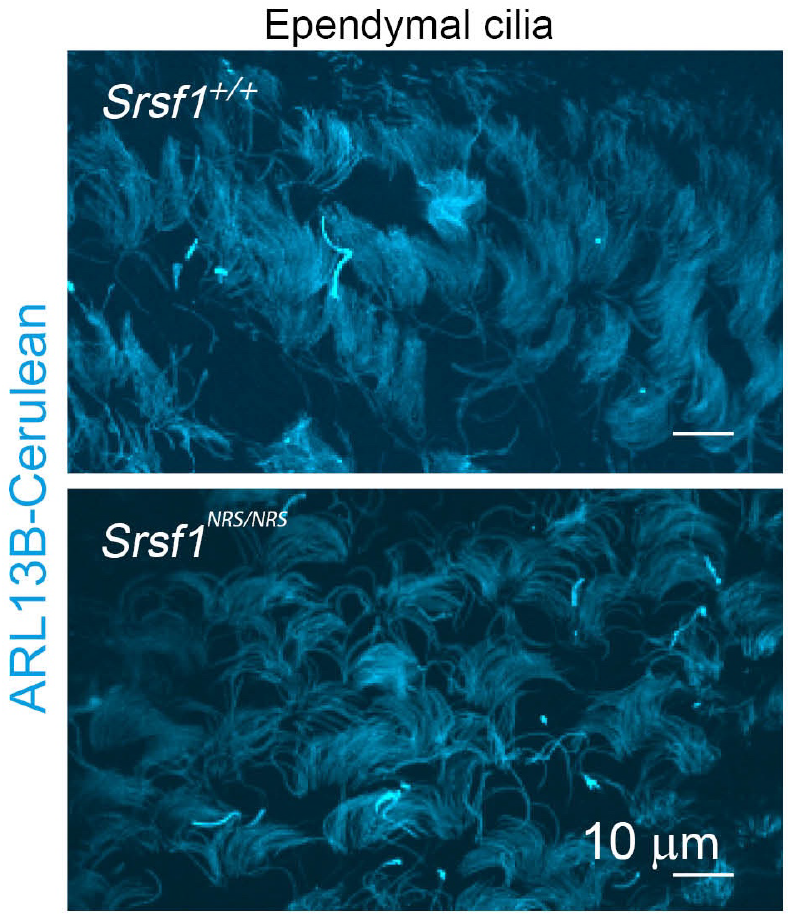
Live images of ARL13B-Cerulean cilia of ependymal cells from the lateral ventricles showing comparable numbers and distribution of ciliary bundles between *Srsf1^NRS/NRS^* and *Srsf1^+/+^* littermates.

**Figure 4-figure supplement 1.**
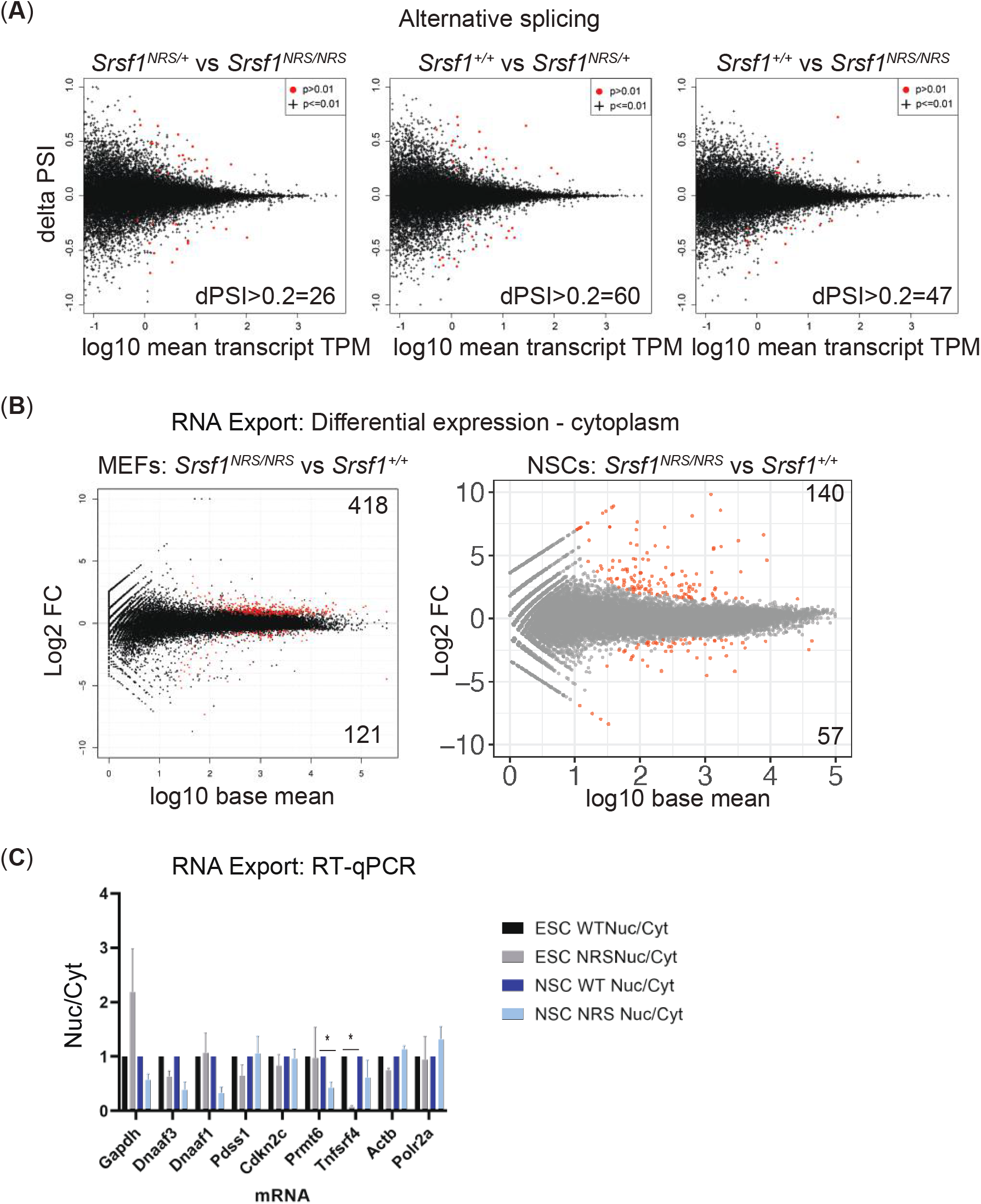
The lack of cytoplasmic SRSF1 does not affect pre-mRNA splicing or mRNA export. (**A**) delta PSI scatter plots for knock-in MEFs. The number of changes (ΔPSI=>0.5, p=<0.01, TPM=>0.5) from indicated pairwise comparisons of splicing changes between each genotype of SRSF1 knock-in mice is depicted. (**B**) Plot of the Log2 fold change of the cytoplasmic abundance of mRNAs between wild type and SRSF1-NRS-expressing MEFs and NSCs. (**C**) The nuclear-cytoplasmic ratio of individual mRNAs in wild-type or SRSF1-NRS expressing cells was quantified by RT-qPCR. The data is an average of three independent experiments, each bar represents an average and a standard error.

**Figure 4-figure supplement 2.**
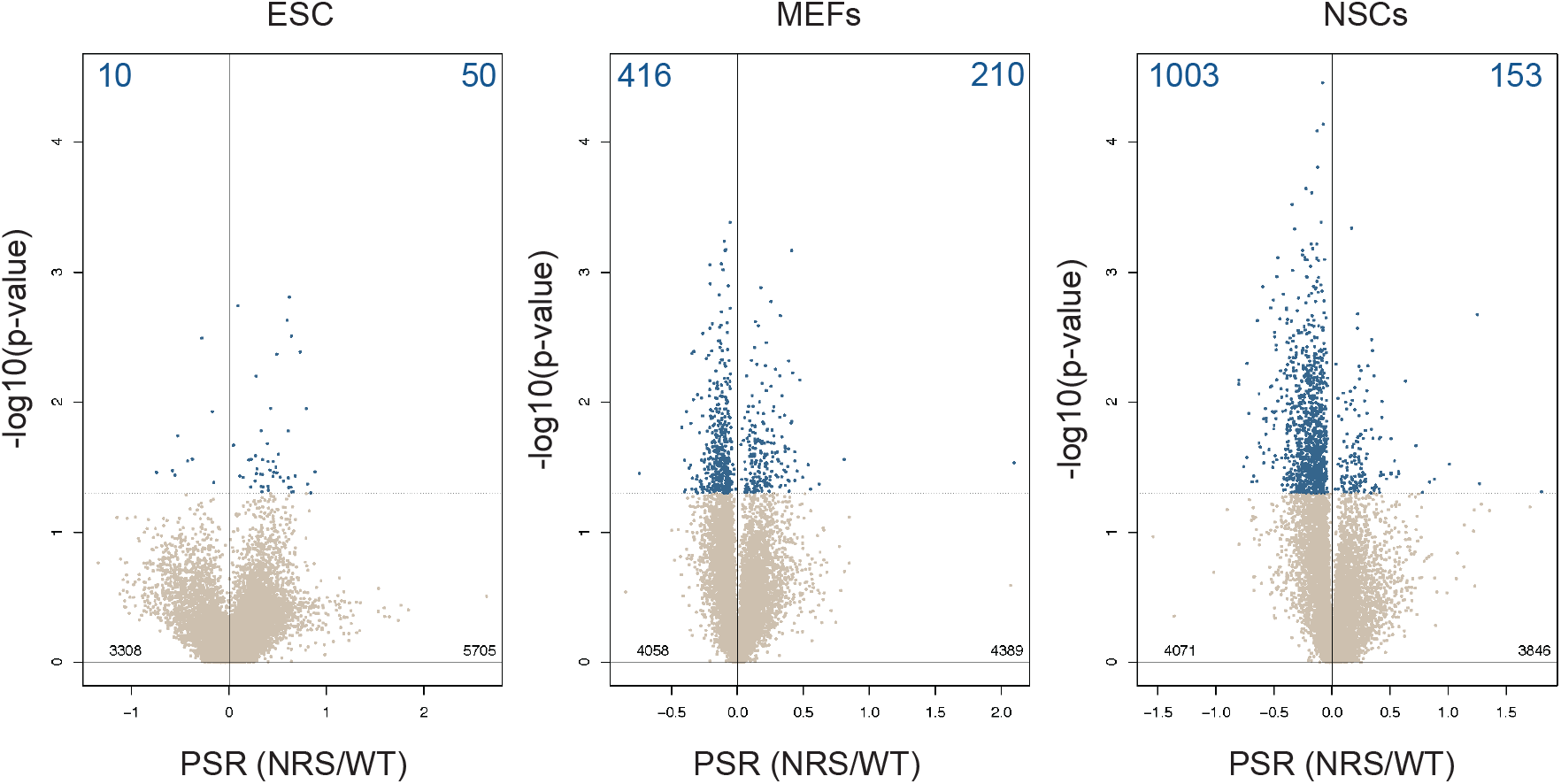
Lack of cytoplasmic SRSF1 induces translational changes not restricted to lineage specific transcript. Scatter plots showing the distribution of genes expressed in each cell line according to their polysome shift ratio (PSR). Blue dots indicate significant changes (FDR<0.05). Number of significant changes is indicated in the top corners of each plot.

**Figure 4-figure supplement 3.**
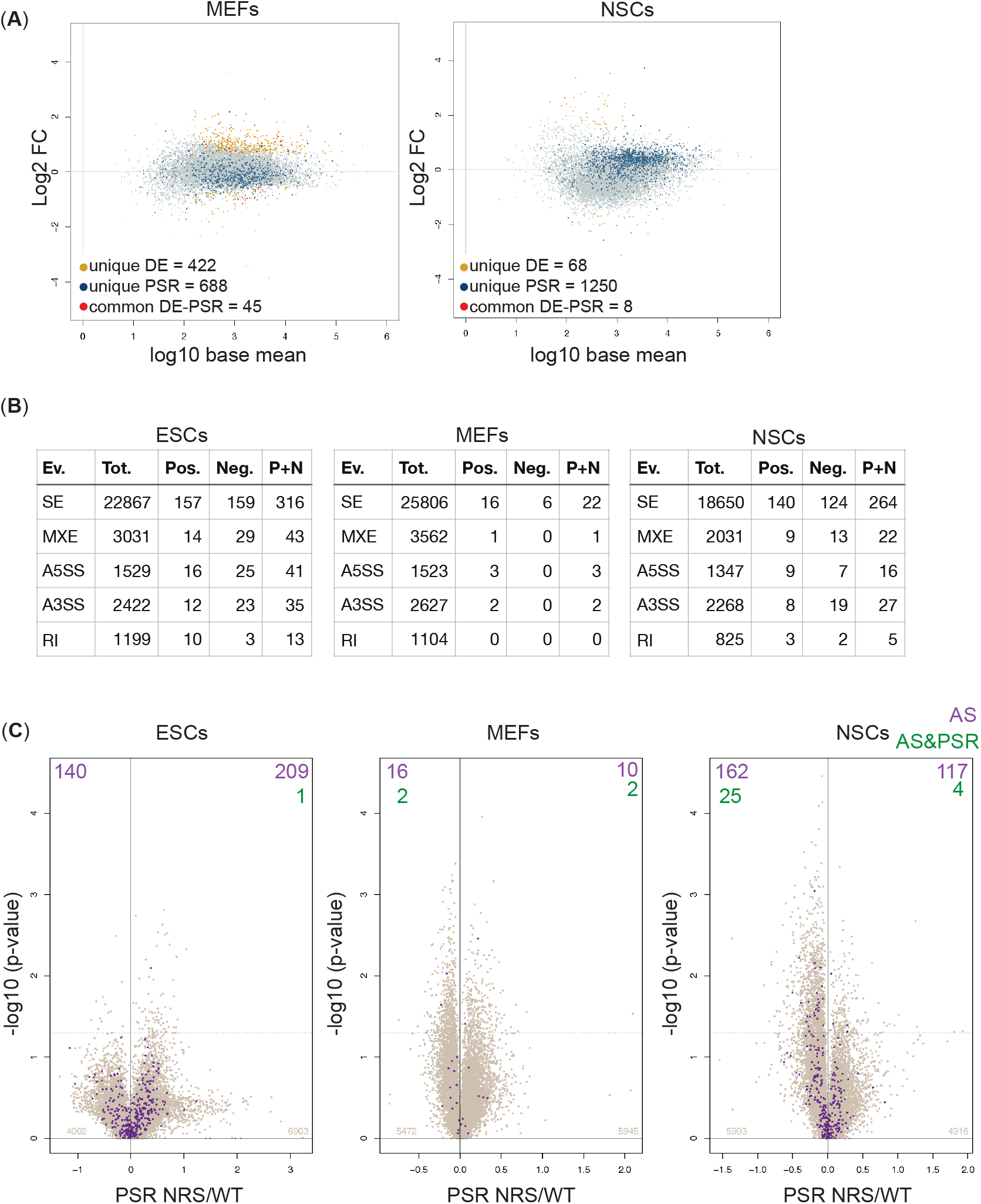
PSR values are independent of alternative splicing and gene expression changes in cytoplasm. (**A**) Plot of the Log2 fold change of the cytoplasmic abundance of mRNAs between *Srsf1^+/+^* and *Srsf1^NRS/NRS^* cells. Yellow, dark blue and red dots indicate significant changes in expression (differential expression, DE), in polysome shift ratio (PSR) or both expression and PSR (DE-PSR), respectively. (**B**) Number of alternative splicing changes detected between *Srsf1^+/+^* and *Srsf1^NRS/NRS^* cells. Ev, type of splicing event: SE, cassette exons; MXE, microexons; A5SS, alternative 5’ splice site; A3SS, alternative 3’ splice site; RI, intron retention. Tot. represents a total number of splicing events, Pos. and Neg. represent those splicing events that increase and decrease in *Srsf1^NRS/NRS^* cells in comparison to *Srsf1^+/+^* cells, respectively. (**C**) Scatter plots showing the distribution of genes expressed in all cell lines according to PSR. Purple dots indicate identified splicing changes. Number of significant splicing changes and significant splicing changes observed for genes with significant PSRs is indicated in the top corners of each plot (AS and AS&PSR, respectively).

**Figure 5-figure supplement 1.**
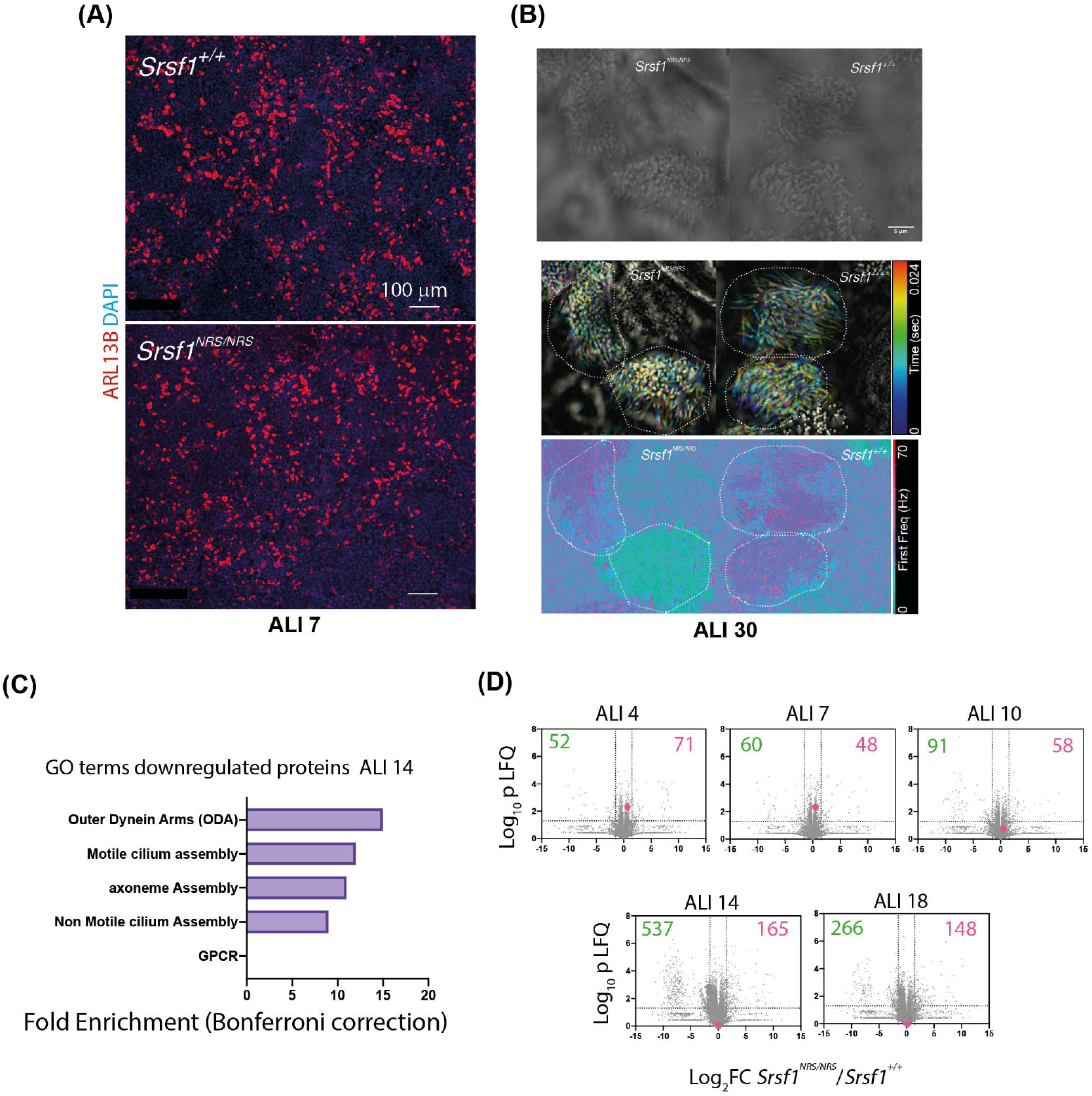
Nuclear sequestration of SRSF1 leads to alterations in total proteomes consistent with defects in cilia motility observed during mTEC differentiation. (**A**) Immunofluorescence of mTECs at ALI7 showing comparable number of ciliated bundles (ARL13B stained) between *Srsf1^+/+^* and *Srsf^1NRS/NRS^* cultures. (**B**) High speed imaging of mature mTECs (ALI30) show different waveforms in *Srsf1^NRS/NRS^* cilia. The still images show a single time frame of the bundles analyzed below. Middle: Color-coded time projections applying the subtract background function in ImageJ. While neighbored cilia on *Srsf1^+/+^* cells align their beat, cilia on *Srsf1^NRS/NRS^* cells beat in a less coordinated. Bottom: Frequency of the highest peak in the power spectrum derived by fast Fourier transformation of the intensity time-course at each pixel, revealing the ciliary beat frequency of individual cells. (**C**) Functional GO terms enriched in downregulated proteins at ALI14 from *Srsf1^NRS/NRS^* mTEC cultures (right panel). (**D**) Volcano plots at indicated timepoints show proteins miss-regulated in *Srsf1^NRS/NRS^* relative to *Srsf1^+/+^* mTEC cultures. Number of proteins altered are shown in each quadrant. Red dots represent SRSF1 peptides.

## References

Al Jord A, Shihavuddin A, D’Aout RS, Faucourt M, Genovesio A, Karaiskou A, Sobczak-Thépot J, Spassky N, Meunier A. 2017. Calibrated mitotic oscillator drives motile ciliogenesis. Science 358:803–806. doi:10.1126/science.aan8311

Anczukow O, Krainer AR. 2016. Splicing-factor alterations in cancers. RNA 22:1285–301. doi:10.1261/rna.057919.116

Anczuków O, Rosenberg AZ, Akerman M, Das S, Zhan L, Karni R, Muthuswamy SK, Krainer AR. 2012. The splicing factor SRSF1 regulates apoptosis and proliferation to promote mammary epithelial cell transformation. Nat Struct Mol Biol 19:220–8. doi:10.1038/nsmb.2207

Baralle FE, Giudice J. 2017. Alternative splicing as a regulator of development and tissue identity. Nat Rev Mol Cell Biol 18:437–451. doi:10.1038/nrm.2017.27

Bedard KM, Daijogo S, Semler BL. 2007. A nucleo-cytoplasmic SR protein functions in viral IRES-mediated translation initiation. EMBO J 26:459–67. doi:10.1038/sj.emboj.7601494

Blair JD, Hockemeyer D, Doudna JA, Bateup HS, Floor SN. 2017. Widespread Translational Remodeling during Human Neuronal Differentiation. Cell Rep 21:2005–2016. doi:10.1016/j.celrep.2017.10.095

Bonnal SC, López-Oreja I, Valcárcel J. 2020. Roles and mechanisms of alternative splicing in cancer — implications for care. Nat Rev Clin Oncol. doi:10.1038/s41571-020-0350-x

Botti V, McNicoll FF, Steiner MC, Richter FM, Solovyeva A, Wegener M, Schwich OD, Poser I, Zarnack K, Wittig I, Neugebauer KM, Müller-McNicoll M, üller-McNicoll M, Müller-McNicoll M. 2017. Cellular differentiation state modulates the mRNA export activity of SR proteins. J Cell Biol 216:1993–2009. doi:10.1083/jcb.201610051

Busch A, Hertel KJ. 2012. Evolution of SR protein and hnRNP splicing regulatory factors. Wiley Interdiscip Rev RNA 3:1–12. doi:10.1002/wrna.100

Cáceres JF, Misteli T, Screaton GR, Spector DL, Krainer AR. 1997. Role of the modular domains of SR proteins in subnuclear localization and alternative splicing specificity. J Cell Biol 138:225–38. doi:10.1083/jcb.138.2.225.

Cáceres JF, Screaton GR, Krainer AR. 1998. A specific subset of SR proteins shuttles continuously between the nucleus and the cytoplasm. Genes Dev 12:55–66. doi:10.1101/gad.12.1.55.

Cazalla D, Zhu J, Manche L, Huber E, Krainer AR, Caceres JF. 2002. Nuclear Export and Retention Signals in the RS Domain of SR Proteins. Mol Cell Biol 22:6871–6882. doi:10.1128/MCB.22.19.6871

Cléry A, Sinha R, Anczuków O, Corrionero A, Moursy A, Daubner GM, Valcárcel J, Krainer AR, Allain FH-T. 2013. Isolated pseudo-RNA-recognition motifs of SR proteins can regulate splicing using a noncanonical mode of RNA recognition. Proc Natl Acad Sci U S A 110:E2802–11. doi:10.1073/pnas.1303445110

Cong L, Ran FA, Cox D, Lin S, Barretto R, Habib N, Hsu PD, Wu X, Jiang W, Marraffini LA, Zhang F. 2013. Multiplex genome engineering using CRISPR/Cas systems. Science 339:819–23. doi:10.1126/science.1231143

Cowper AE, Cáceres JF, Mayeda A, Screaton GR. 2001. Serine-arginine (SR) protein-like factors that antagonize authentic SR proteins and regulate alternative splicing. J Biol Chem 276:48908–14. doi:10.1074/jbc.M103967200

Crichton JH, Playfoot CJ, MacLennan M, Read D, Cooke HJ, Adams IR. 2017. Tex19.1 promotes Spo11-dependent meiotic recombination in mouse spermatocytes. PLOS Genet 13:e1006904. doi:10.1371/journal.pgen.1006904

Czechanski A, Byers C, Greenstein I, Schrode N, Donahue LR, Hadjantonakis AK, Reinholdt LG. 2014. Derivation and characterization of mouse embryonic stem cells from permissive and nonpermissive strains. Nat Protoc 9:559–574. doi:10.1038/nprot.2014.030

Dobin A, Davis CA, Schlesinger F, Drenkow J, Zaleski C, Jha S, Batut P, Chaisson M, Gingeras TR. 2013. STAR: Ultrafast universal RNA-seq aligner. Bioinformatics 29:15–21. doi:10.1093/bioinformatics/bts635

Fingerhut JM, Yamashita YM. 2020. mRNA localization mediates maturation of cytoplasmic cilia in Drosophila spermatogenesis. J Cell Biol 219:e202003084. doi:10.1083/jcb.202003084

Ford MJ, Yeyati PL, Mali GR, Keighren MA, Waddell SH, Mjoseng HK, Douglas AT, Hall EA, Sakaue-Sawano A, Miyawaki A, Meehan RR, Boulter L, Jackson IJ, Mill P, Mort RL. 2018. A Cell/Cilia Cycle Biosensor for Single-Cell Kinetics Reveals Persistence of Cilia after G1/S Transition Is a General Property in Cells and Mice. Dev Cell 47:509–523.e5. doi:10.1016/j.devcel.2018.10.027

Fu X-D, Ares M. 2014. Context-dependent control of alternative splicing by RNA-binding proteins. Nat Rev Genet 15:689–701. doi:10.1038/nrg3778

Fu Y, Huang B, Shi Z, Han J, Wang Y, Huangfu J, Wu W. 2013. SRSF1 and SRSF9 RNA binding proteins promote Wnt signalling-mediated tumorigenesis by enhancing β-catenin biosynthesis. EMBO Mol Med 5:737–50. doi:10.1002/emmm.201202218

Gomperts BN, Gong-Cooper X, Hackett BP. 2004. Foxj1 regulates basal body anchoring to the cytoskeleton of ciliated pulmonary epithelial cells. J Cell Sci 117:1329–1337. doi:10.1242/jcs.00978

Gui JF, Lane WS, Fu XD. 1994. A serine kinase regulates intracellular localization of splicing factors in the cell cycle. Nature 369:678–82. doi:10.1038/369678a0

Hanamura A, Cáceres JF, Mayeda A, Franza BR, Krainer AR. 1998. Regulated tissue-specific expression of antagonistic pre-mRNA splicing factors. RNA 4:430–44.

Hansen J, Rassmann S, Jikeli J, Wachten D. 2018. SpermQ–A Simple Analysis Software to Comprehensively Study Flagellar Beating and Sperm Steering. Cells 8:10. doi:10.3390/cells8010010

Hargous Y, Hautbergue GM, Tintaru AM, Skrisovska L, Golovanov AP, Stevenin J, Lian L-Y, Wilson SA, Allain FH-T. 2006. Molecular basis of RNA recognition and TAP binding by the SR proteins SRp20 and 9G8. EMBO J 25:5126–37. doi:10.1038/sj.emboj.7601385

Howard JM, Sanford JR. 2015. The RNAissance family: SR proteins as multifaceted regulators of gene expression. Wiley Interdiscip Rev RNA 6:93–110. doi:10.1002/wrna.1260

Huang Y, Gattoni R, Stévenin J, Steitz JA. 2003. SR splicing factors serve as adapter proteins for TAP-dependent mRNA export. Mol Cell 11:837–43. doi:10.1016/s1097-2765(03)00089-3

Huang Y, Steitz JA. 2001. Splicing factors SRp20 and 9G8 promote the nucleocytoplasmic export of mRNA. Mol Cell 7:899–905.

Huizar RL, Lee C, Boulgakov AA, Horani A, Tu F, Marcotte EM, Brody SL, Wallingford JB. 2018. A liquid-like organelle at the root of motile ciliopathy. Elife 7:e38497. doi:10.7554/eLife.38497

Ibañez-Tallon I, Pagenstecher A, Fliegauf M, Olbrich H, Kispert A, Ketelsen UP, North A, Heintz N, Omran H. 2004. Dysfunction of axonemal dynein heavy chain Mdnah5 inhibits ependymanl flow and reveals a novel mechanism for hydrocephalus formation. Hum Mol Genet 13:2133–2141. doi:10.1093/hmg/ddh219

Jain R, Pan J, Driscoll JA, Wisner JW, Huang T, Gunsten SP, You Y, Brody SL. 2010. Temporal relationship between primary and motile ciliogenesis in airway epithelial cells. Am J Respir Cell Mol Biol 43:731–739. doi:10.1165/rcmb.2009-0328OC

Joyner AL. 2000. Gene targeting: a practical approach. Oxford University Press.

Karni R, de Stanchina E, Lowe SW, Sinha R, Mu D, Krainer AR. 2007. The gene encoding the splicing factor SF2/ASF is a proto-oncogene. Nat Struct Mol Biol 14:185–93. doi:10.1038/nsmb1209

Kataoka N, Bachorik JL, Dreyfuss G. 1999. Transportin-SR, a nuclear import receptor for SR proteins. J Cell Biol 145:1145–52. doi:10.1083/jcb.145.6.1145

Katsuyama T, Li H, Comte D, Tsokos GC, Moulton VR. 2019. Splicing factor SRSF1 controls T cell hyperactivity and systemic autoimmunity. J Clin Invest 129:5411–23. doi:10.1172/JCI127949

Lai MC, Lin RI, Huang SY, Tsai CW, Tarn WY. 2000. A human importin-beta family protein, transportin-SR2, interacts with the phosphorylated RS domain of SR proteins. J Biol Chem 275:7950–7. doi:10.1074/jbc.275.11.7950

Langdon EM, Qiu Y, Niaki AG, McLaughlin GA, Weidmann CA, Gerbich TM, Smith JA, Crutchley JM, Termini CM, Weeks KM, Myong S, Gladfelter AS. 2018. mRNA structure determines specificity of a polyQ-driven phase separation. Science 360:922–927. doi:10.1126/science.aar7432

Lewis M, Stracker TH. 2020. Transcriptional regulation of multiciliated cell differentiation. Semin Cell Dev Biol. doi:10.1016/j.semcdb.2020.04.007

Liao Y, Wang J, Jaehnig EJ, Shi Z, Zhang B. 2019. WebGestalt 2019: gene set analysis toolkit with revamped UIs and APIs. Nucleic Acids Res 47:W199–W205. doi:10.1093/nar/gkz401

Lin S, Xiao R, Sun P, Xu X, Fu X-D. 2005. Dephosphorylation-dependent sorting of SR splicing factors during mRNP maturation. Mol Cell 20:413–25. doi:10.1016/j.molcel.2005.09.015

Liu Y, Luo Y, Shen L, Guo R, Zhan Z, Yuan N, Sha R, Qian W, Wang Z, Xie Z, Wu W, Feng Y. 2020. Splicing Factor SRSF1 Is Essential for Satellite Cell Proliferation and Postnatal Maturation of Neuromuscular Junctions in Mice. Stem Cell Reports. doi:10.1016/j.stemcr.2020.08.004

Lobo J, Zariwala MA, Noone PG. 2015. Primary ciliary dyskinesia. Semin Respir Crit Care Med 36:169–179. doi:10.1055/s-0035-1546748

Loges NT, Olbrich H, Becker-Heck A, Häffner K, Heer A, Reinhard C, Schmidts M, Kispert A, Zariwala MA, Leigh MW, Knowles MR, Zentgraf H, Seithe H, Nürnberg G, Nürnberg P, Reinhardt R, Omran H. 2009. Deletions and Point Mutations of LRRC50 Cause Primary Ciliary Dyskinesia Due to Dynein Arm Defects. Am J Hum Genet 85:883–889. doi:10.1016/j.ajhg.2009.10.018

Loges NT, Olbrich H, Fenske L, Mussaffi H, Horvath J, Fliegauf M, Kuhl H, Baktai G, Peterffy E, Chodhari R, Chung EMK, Rutman A, O’Callaghan C, Blau H, Tiszlavicz L, Voelkel K, Witt M, Zietkiewicz E, Neesen J, Reinhardt R, Mitchison HM, Omran H. 2008. DNAI2 Mutations Cause Primary Ciliary Dyskinesia with Defects in the Outer Dynein Arm. Am J Hum Genet 83:547–558. doi:10.1016/j.ajhg.2008.10.001

Long JC, Caceres JF. 2009. The SR protein family of splicing factors: master regulators of gene expression. Biochem J 417:15–27. doi:10.1042/BJ20081501

Love MI, Huber W, Anders S. 2014. Moderated estimation of fold change and dispersion for RNA-seq data with DESeq2. Genome Biol 15:550. doi:10.1186/s13059-014-0550-8

Maharana S, Wang J, Papadopoulos DK, Richter D, Pozniakovsky A, Poser I, Bickle M, Rizk S, Guillén-Boixet J, Franzmann TM, Jahnel M, Marrone L, Chang YT, Sterneckert J, Tomancak P, Hyman AA, Alberti S. 2018. RNA buffers the phase separation behavior of prion-like RNA binding proteins. Science 360:918–921. doi:10.1126/science.aar7366

Maslon MM, Braunschweig U, Aitken S, Mann AR, Kilanowski F, Hunter CJ, Blencowe BJ, Kornblihtt AR, Adams IR, Cáceres JF. 2019. A slow transcription rate causes embryonic lethality and perturbs kinetic coupling of neuronal genes. EMBO J 38:e101244. doi:10.15252/embj.2018101244

Maslon MM, Heras SR, Bellora N, Eyras E, Cáceres JF. 2014. The translational landscape of the splicing factor SRSF1 and its role in mitosis. Elife 3:e02028. doi:10.7554/eLife.02028

McAllister JP. 2012. Pathophysiology of congenital and neonatal hydrocephalus. Semin Fetal Neonatal Med. doi:10.1016/j.siny.2012.06.004

Meunier A, Azimzadeh J. 2016. Multiciliated cells in animals. Cold Spring Harb Perspect Biol 8:a028233. doi:10.1101/cshperspect.a028233

Michlewski G, Sanford JR, Cáceres JF. 2008. The splicing factor SF2/ASF regulates translation initiation by enhancing phosphorylation of 4E-BP1. Mol Cell 30:179–89. doi:10.1016/j.molcel.2008.03.013

Mitchison HM, Schmidts M, Loges NT, Freshour J, Dritsoula A, Hirst RA, O’Callaghan C, Blau H, Al Dabbagh M, Olbrich H, Beales PL, Yagi T, Mussaffi H, Chung EMK, Omran H, Mitchell DR. 2012. Mutations in axonemal dynein assembly factor DNAAF3 cause primary ciliary dyskinesia. Nat Genet 44:381–389. doi:10.1038/ng.1106

Möröy T, Heyd F. 2007. The impact of alternative splicing in vivo: Mouse models show the way. RNA. doi:10.1261/rna.554607

Müller-McNicoll M, Botti V, de Jesus Domingues AM, Brandl H, Schwich OD, Steiner MC, Curk T, Poser I, Zarnack K, Neugebauer KM. 2016. SR proteins are NXF1 adaptors that link alternative RNA processing to mRNA export. Genes Dev 30:553–66. doi:10.1101/gad.276477.115

Naftelberg S, Schor IE, Ast G, Kornblihtt AR. 2015. Regulation of Alternative Splicing Through Coupling with Transcription and Chromatin Structure. Annu Rev Biochem 84:165–198. doi:10.1146/annurev-biochem-060614-034242

Nilsen TW, Graveley BR. 2010. Expansion of the eukaryotic proteome by alternative splicing. Nature 463:457–63. doi:10.1038/nature08909

Olstad EW, Ringers C, Hansen JN, Wens A, Brandt C, Wachten D, Yaksi E, Jurisch-Yaksi N. 2019. Ciliary Beating Compartmentalizes Cerebrospinal Fluid Flow in the Brain and Regulates Ventricular Development. Curr Biol 29:229–241.e6. doi:10.1016/j.cub.2018.11.059

Oltean A, Schaffer AJ, Bayly P V., Brody SL. 2018. Quantifying ciliary dynamics during assembly reveals stepwise waveform maturation in airway cells. Am J Respir Cell Mol Biol 59:511–522. doi:10.1165/rcmb.2017-0436OC

Park JW, Tokheim C, Shen S, Xing Y. 2013. Identifying differential alternative splicing events from RNA sequencing data using RNASeq-MATS. Methods Mol Biol 1038:171–179. doi:10.1007/978-1-62703-514-9_10

Patro R, Duggal G, Love MI, Irizarry RA, Kingsford C. 2017. Salmon provides fast and bias-aware quantification of transcript expression. Nat Methods 14:417–419. doi:10.1038/nmeth.4197

Paz S, Ritchie A, Mauer C, Caputi M. 2020. The RNA binding protein SRSF1 is a master switch of gene expression and regulation in the immune system. Cytokine Growth Factor Rev. doi:10.1016/j.cytogfr.2020.10.008

Piñol-Roma S, Dreyfuss G. 1992. Shuttling of pre-mRNA binding proteins between nucleus and cytoplasm. Nature 355:730–2. doi:10.1038/355730a0

Prasad J, Colwill K, Pawson T, Manley JL. 1999. The protein kinase Clk/Sty directly modulates SR protein activity: both hyper- and hypophosphorylation inhibit splicing. Mol Cell Biol 19:6991–7000. doi:10.1128/mcb.19.10.6991.

Rappsilber J, Ishihama Y, Mann M. 2003. Stop And Go Extraction tips for matrix-assisted laser desorption/ionization, nanoelectrospray, and LC/MS sample pretreatment in proteomics. Anal Chem 75:663–670. doi:10.1021/ac026117i

Reiter JF, Leroux MR. 2017. Genes and molecular pathways underpinning ciliopathies. Nat Rev Mol Cell Biol. doi:10.1038/nrm.2017.60

Saldi T, Cortazar MA, Sheridan RM, Bentley DL. 2016. Coupling of RNA polymerase II transcription elongation with pre-mRNA splicing. J Mol Biol 428:2623–35. doi:10.1016/j.jmb.2016.04.017

Sanford JR, Ellis JD, Cazalla D, Cáceres JF. 2005. Reversible phosphorylation differentially affects nuclear and cytoplasmic functions of splicing factor 2/alternative splicing factor. Proc Natl Acad Sci U S A 102:15042–7. doi:10.1073/pnas.0507827102

Sanford JR, Gray NK, Beckmann K, Cáceres JF. 2004. A novel role for shuttling SR proteins in mRNA translation. Genes Dev 18:755–768. doi:10.1101/gad.286404

Sapra AK, Ankö M-L, Grishina I, Lorenz M, Pabis M, Poser I, Rollins J, Weiland E-M, Neugebauer KM. 2009. SR protein family members display diverse activities in the formation of nascent and mature mRNPs in vivo. Mol Cell 34:179–90. doi:10.1016/j.molcel.2009.02.031

Schindelin J, Arganda-Carreras I, Frise E, Kaynig V, Longair M, Pietzsch T, Preibisch S, Rueden C, Saalfeld S, Schmid B, Tinevez JY, White DJ, Hartenstein V, Eliceiri K, Tomancak P, Cardona A. 2012. Fiji: An open-source platform for biological-image analysis. Nat Methods. doi:10.1038/nmeth.2019

Sun S, Zhang Z, Sinha R, Karni R, Krainer AR. 2010. SF2/ASF autoregulation involves multiple layers of post-transcriptional and translational control. Nat Struct Mol Biol 17:306–12. doi:10.1038/nsmb.1750

Swartz JE, Bor Y-C, Misawa Y, Rekosh D, Hammarskjold M-L. 2007. The shuttling SR protein 9G8 plays a role in translation of unspliced mRNA containing a constitutive transport element. J Biol Chem 282:19844–53. doi:10.1074/jbc.M701660200

Tanaka H, Iguchi N, Toyama Y, Kitamura K, Takahashi T, Kaseda K, Maekawa M, Nishimune Y. 2004. Mice Deficient in the Axonemal Protein Tektin-t Exhibit Male Infertility and Immotile-Cilium Syndrome Due to Impaired Inner Arm Dynein Function. Mol Cell Biol 24:7958–7964. doi:10.1128/mcb.24.18.7958-7964.2004

Trincado JL, Entizne JC, Hysenaj G, Singh B, Skalic M, Elliott DJ, Eyras E. 2018. SUPPA2: fast, accurate, and uncertainty-aware differential splicing analysis across multiple conditions. Genome Biol 19:40. doi:10.1186/s13059-018-1417-1

Twyffels L, Gueydan C, Kruys V. 2011. Shuttling SR proteins: more than splicing factors. FEBS J 278:3246–55. doi:10.1111/j.1742-4658.2011.08274.x

Ule J, Blencowe BJ. 2019. Alternative Splicing Regulatory Networks: Functions, Mechanisms, and Evolution. Mol Cell. doi:10.1016/j.molcel.2019.09.017

Vladar EK, Brody SL. 2013. Analysis of ciliogenesis in primary culture mouse tracheal epithelial cellsMethods in Enzymology. Academic Press Inc. pp. 285–309. doi:10.1016/B978-0-12-397944-5.00014-6

Wang J, Takagaki Y, Manley JL. 1996. Targeted disruption of an essential vertebrate gene: ASF/SF2 is required for cell viability. Genes Dev 10:2588–99. doi:10.1101/gad.10.20.2588

Wegener M, Müller-McNicoll M. 2019. View from an mRNP: The Roles of SR Proteins in Assembly, Maturation and Turnover. Adv Exp Med Biol 1203:83–112. doi:10.1007/978-3-030-31434-7_3

Wirschell M, Olbrich H, Werner C, Tritschler D, Bower R, Sale WS, Loges NT, Pennekamp P, Lindberg S, Stenram U, Carlén B, Horak E, Köhler G, Nürnberg P, Nürnberg G, Porter ME, Omran H. 2013. The nexin-dynein regulatory complex subunit DRC1 is essential for motile cilia function in algae and humans. Nat Genet 45:262–268. doi:10.1038/ng.2533

Xu X, Yang D, Ding J-H, Wang W, Chu P-H, Dalton ND, Wang H-Y, Bermingham JR, Ye Z, Liu F, Rosenfeld MG, Manley JL, Ross J, Chen J, Xiao R-P, Cheng H, Fu X-D. 2005. ASF/SF2-regulated CaMKIIdelta alternative splicing temporally reprograms excitation-contraction coupling in cardiac muscle. Cell 120:59–72. doi:10.1016/j.cell.2004.11.036

Zhang J, Manley JL. 2013. Misregulation of pre-mRNA alternative splicing in cancer. Cancer Discov. doi:10.1158/2159-8290.CD-13-0253

Zhang YJ, O’Neal WK, Randell SH, Blackburn K, Moyer MB, Boucher RC, Ostrowski LE. 2002. Identification of dynein heavy chain 7 as an inner arm component of human cilia that is synthesized but not assembled in a case of primary ciliary dyskinesia. J Biol Chem 277:17906–17915. doi:10.1074/jbc.M200348200

Zhou Z, Fu X-D. 2013. Regulation of splicing by SR proteins and SR protein-specific kinases. Chromosoma 122:191–207. doi:10.1007/s00412-013-0407-z

Zhu J, Mayeda A, Krainer AR. 2001. Exon identity established through differential antagonism between exonic splicing silencer-bound hnRNP A1 and enhancer-bound SR proteins. Mol Cell 8:1351–61. doi:doi:10.1016/s1097-2765(01)00409-9.

